# Prenatal Zika Virus Exposure Disrupts Social-Emotional Development and Cortical Visual Function in Infant Macaques

**DOI:** 10.1101/2025.06.09.658678

**Authors:** Karla K. Ausderau, Ben Boerigter, Elaina R. Razo, Jake Gutkes, Nicholas P. Krabbe, Ann M. Mitzey, Shannon Walsh, Viktorie Menna, John R. Drew, Sabrina Kabakov, Finn Eckes, Rachel V. Spanton, Anika Shah, Angelica Sun, Alex Katz, Charlene Kim, Amy Hartman, Andrea M. Weiler, Carol Rasmussen, Micheal Nork, Puja Basu, Heather A. Simmons, James Ver Hoeve, Saverio Capuano, Thomas C. Friedrich, Emma L. Mohr

## Abstract

Prenatal exposure to Zika virus (ZIKV) results in a spectrum of outcomes, ranging from severe birth defects and early childhood developmental delays to no apparent deficits. However, the mechanisms underlying these outcomes, particularly in the context of different maternal infection conditions, remain poorly defined. In this study, we employed a translational rhesus macaque model of prenatal ZIKV infection to evaluate longitudinal visual and auditory development from 1 to 12 months of age and characterize neurodevelopmental outcomes at 12 months. Pregnant macaques, either flavivirus-naive or with prior dengue virus (DENV) exposure, were inoculated in the first trimester (∼30 or 45 gestational days) with an Asian or African lineage of ZIKV or received a saline injection as controls. Maternal plasma viremia duration, viral RNA burden at the maternal-fetal interface, and neutralizing antibody titers did not differ between inoculation cohorts or controls. At 12 months, ZIKV-exposed infants demonstrated altered maternal attachment behaviors and reduced inhibition when approaching sensory stimuli compared to controls. Visual pathway function, assessed by electrophysiology, was significantly impaired at 3 months but normalized by 12 months. Hearing loss was more common among ZIKV-exposed infants, although not statistically significant. Developmental outcomes were associated with prenatal ZIKV exposure itself, independent of viremia duration, neutralizing antibody titer, viral lineage, or maternal DENV immunity, and were not mediated by visual impairment or hearing loss. These findings highlight that any prenatal ZIKV exposure can disrupt early neurobehavioral development and visual function, underscoring the need for prevention strategies focused on maternal infection and early intervention.

## INTRODUCTION

Prenatal Zika virus (ZIKV) exposure produces a broad range of infant outcomes, including birth defects, neurodevelopmental delays, hearing loss, and visual impairment. ZIKV continues to cause periodic outbreaks in endemic regions, while emerging transmission is being detected in previously unaffected areas (*1*, *2*). Notably, nearly 30% of affected human infants are asymptomatic at birth but develop neurodevelopmental deficits during early childhood (*3*, *4*).

These deficits span language, cognitive, motor, hearing, and visual domains (*4–8*), alongside impairments in mobility, communication, and social cognition (*3*, *5*). A smaller proportion (∼6%) present at birth with congenital Zika syndrome, a constellation of findings including microcephaly, brain and ocular anomalies, congenital joint contractures, and neurologic sequelae such as sensorineural hearing loss (*9*, *10*). Prenatal ZIKV infection also impacts vision (*11*) and may impact hearing (*12*, *13*), and early disruptions in vision and hearing are well recognized to profoundly influence language acquisition, cognition, and socio-emotional development (*14*, *15*). Yet, critical gaps remain in understanding how maternal ZIKV infection conditions—including viral dynamics and immune responses—shape these developmental trajectories. Elucidating these relationships could enable earlier risk stratification and inform targeted intervention strategies.

Translational rhesus macaque models provide a uniquely powerful system for understanding how maternal ZIKV infection conditions shape developmental outcomes. Compared with murine models, macaques exhibit closer parallels to human placentation, gestational timelines, and fetal neurodevelopmental maturation (*16*, *17*). Rhesus macaques have been firmly established as models for congenital ZIKV infection (*16–20*) and have a longstanding role in modeling human neurodevelopment (*21*). Our preclinical macaque model bridges critical gaps in clinical research by directly assessing ocular and auditory outcomes (*22*) and by demonstrating visual orientation deficits in offspring with prenatal ZIKV exposure and maternal dengue virus (DENV) immunity (*23*). The convergence of neurodevelopmental similarity and an established infection model uniquely positions rhesus macaques to unravel predictors of childhood developmental impairments.

Maternal viral dynamics and humoral immune responses may influence congenital infection outcomes; however, the relationship between these factors and the most common outcomes, namely developmental deficits, remains unclear. Although the hypothesis that prolonged maternal viremia increases the risk of adverse pregnancy and neonatal outcomes is biologically plausible, supporting data are limited. To date, only one cohort study from French Guiana has directly compared outcomes based on maternal viremia duration, demonstrating that maternal plasma viremia persisting for more than 30 days was associated with an increased risk of fetal loss and structural brain abnormalities (*24*). Additional support for this hypothesis is suggested by multiple human case reports (*25–27*). Maternal neutralizing antibody (nAb) responses may also modulate outcomes, as higher nAb titers measured 36-60 days after acute infection were associated with a reduced risk of microcephaly and structural brain abnormalities in one human study (*28*), although neurodevelopmental outcomes were not assessed. Better characterization of maternal viro-immunologic parameters associated with neurodevelopmental deficits could identify potential targets for future early intervention strategies.

Maternal pre-existing immunity to dengue virus (DENV), resulting from infection prior to pregnancy, may influence offspring outcomes following prenatal Zika virus (ZIKV) exposure. In human pediatric cohorts, prior DENV infection has been associated with reduced risk of symptomatic ZIKV disease, suggesting potential cross-protection (*29*). However, experimental data from in vitro systems, ex vivo placental models, and murine studies raise concerns that DENV-specific IgG could enhance ZIKV infection via antibody-dependent enhancement mechanisms (*30*, *31*). In contrast, translational studies in nonpregnant macaques have shown no evidence that pre-existing DENV immunity increases ZIKV titers or disease severity (*32*, *33*).

We previously demonstrated that maternal DENV immunity can worsen early neurodevelopmental outcomes in neonatal macaques exposed to ZIKV in utero (*23*); however, its impact on later developmental outcomes has not been evaluated. This knowledge gap is increasingly important as DENV incidence rises in parts of South America (*34*), and human studies remain inconclusive—some reporting no association between maternal DENV immunity and adverse pregnancy outcomes (*35–37*), while others suggest a protective effect against congenital Zika syndrome (*38*). Defining how prior maternal DENV immunity modulates the effects of prenatal ZIKV exposure on long-term development is essential to optimize early intervention strategies and inform maternal risk assessment.

The timing of maternal Zika virus (ZIKV) infection during pregnancy critically influences offspring outcomes. First-trimester infections are well established to confer a higher risk of birth defects and microcephaly compared to infections occurring later in pregnancy (*9*, *39–41*).

However, the impact of gestational age at exposure on longer-term developmental outcomes is less well-studied. A recent study reported that children with prenatal ZIKV exposure exhibited motor and cognitive delays, with first-trimester exposure associated with an 11.2-fold greater odds of adverse developmental outcomes compared to third-trimester exposure (*7*). Nonetheless, the first trimester encompasses a broad period of early development, and it remains unclear whether the timing within the first trimester further modifies the risk of developmental deficits.

Finally, infection with different ZIKV lineages may impact developmental outcomes. Two primary lineages exist—Asian-American and African. While African-lineage ZIKV has circulated since its initial detection in 1947, its role in congenital infection and subsequent developmental impairments remains poorly characterized in humans. In pregnant immunocompromised mouse models, African-lineage strains have demonstrated greater fetal harm and mortality than Asian-lineage strains (*42*). Similarly, nonhuman primate studies report higher burden of ZIKV RNA within the maternal-fetal interface (*43*) and high rates of fetal demise following African-lineage ZIKV infection during early gestation (*44*) compared to Asian-lineage ZIKV. However, whether these findings extrapolate to humans remains unclear. Defining the impact of African-lineage ZIKV on early childhood development is critical to capturing the full clinical spectrum of congenital ZIKV disease.

To address these gaps, we conducted longitudinal assessments of visual and auditory development during the first year of life and characterized neurodevelopmental outcomes at 12 months in a translational rhesus macaque model of prenatal ZIKV exposure. Using the largest cohort of prenatally ZIKV-exposed infant macaques to date (n = 41), we investigated how maternal ZIKV inoculation, viremia duration, and antibody responses influence infant developmental trajectories. These data offer critical insight into the lasting impact of prenatal ZIKV exposure and the limitations of maternal biomarkers in predicting neurodevelopmental outcomes.

## RESULTS

### Virologic outcomes

Pregnant ZIKV-inoculated animals developed positive plasma ZIKV vRNA loads in the first few days after inoculation (Figure 1A). The duration of positive plasma vRNA loads varied among animals, ranging from 2 to 70 days, but did not differ significantly among dams with different maternal ZIKV inoculation conditions, including inoculation of ZIKV-PR at ∼30 or ∼45 gestational days (gd), history of infection with DENV, or with ZIKV-DAK (Figure 1B, Table S1). Area under the plasma vRNA load curve (AUC) also did not differ between dams in the different maternal ZIKV inoculation conditions (Figure 1C). We also examined the vRNA loads in the maternal interface tissues following Cesarean delivery, specifically examining viral loads separately in the placenta, decidua, and chorionic membrane in biopsies from all the cotyledons in both placentas and found no differences between groups (Figure 1D). When biopsies from the placenta, decidua, and chorionic membrane were combined as a single parameter (% of total biopsies), there was also no difference between dams in the inoculation groups. There were also no differences in the ZIKV-specific neutralizing antibody titers (PRNT90) one month post-inoculation between the different maternal inoculation groups (Figure 1E). We also assessed infants for ZIKV vRNA in plasma and urine, and for ZIKV-specific IgM, as direct markers of fetal infection. None of the infants had detectable ZIKV RNA in plasma (measured longitudinally from birth to 12 months of age) or in urine (measured at birth) (Table S2). None of the infants had detectable ZIKV-specific IgM measured at 0 - 8 days of age (Table S3). Because we detected neither ZIKV RNA nor virus-specific IgM, we refer to these infants as ZIKV-exposed for future discussion, although it is possible that vRNA and/or IgM were present at one time but cleared after the first trimester infection more than 110 days before delivery.

**Fig. 1.**
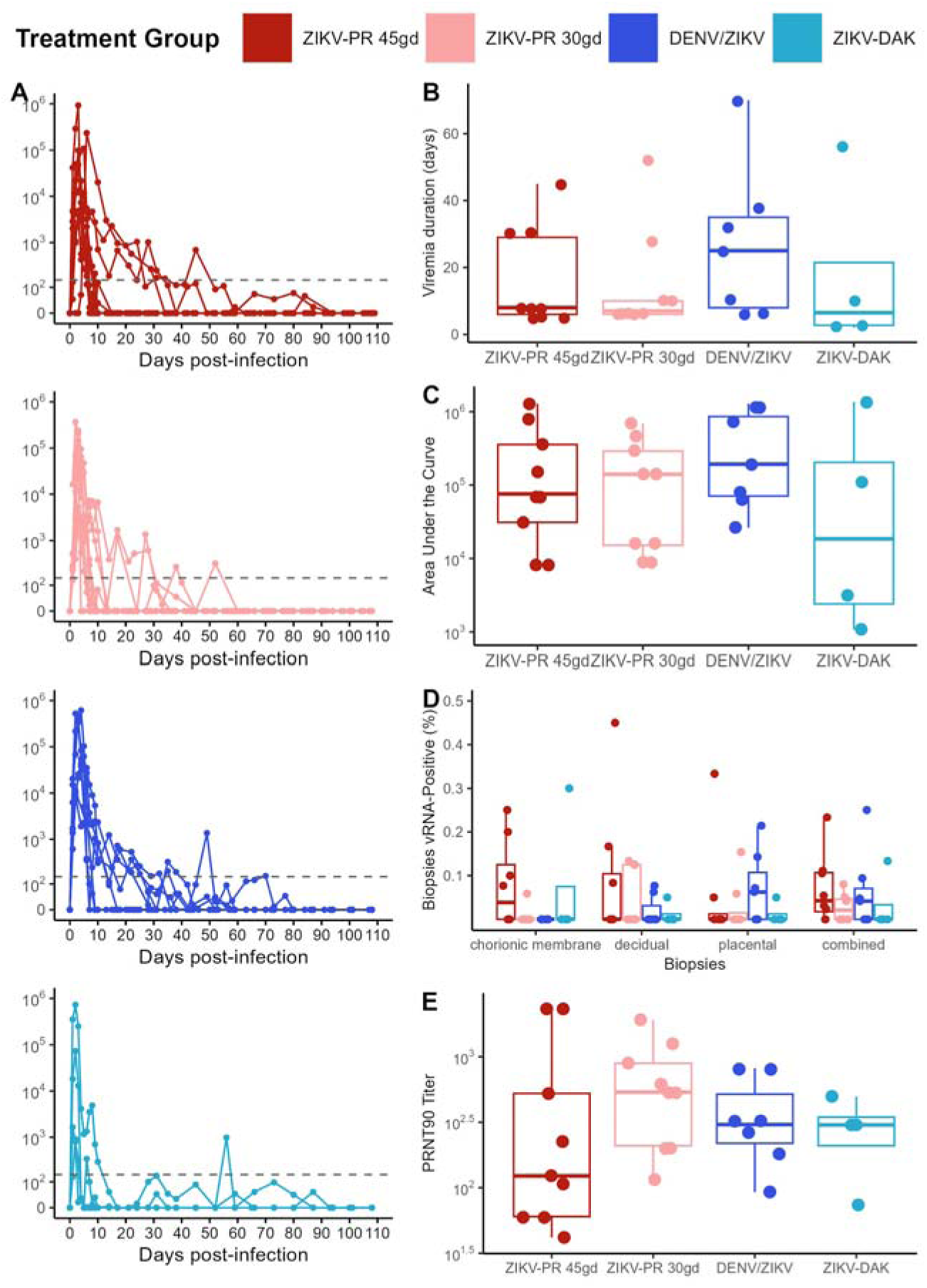
Virologic and neutralizing antibody outcomes. (A) ZIKV vRNA loads in plasma following inoculation in the ZIKV-PR 30gd, ZIKV-PR-45gd, DENV/ZIKV, and ZIKV-DAK groups. The average vRNA load is represented by a solid line and the minimum and maximum vRNA loads by the shaded area. (B) Duration of plasma viremia, defined as the last day of a vRNA load above the limit of detection (150 copies/ml). (C) Area under the plasma viremia curve. (D) The percentage of biopsies from the placenta, decidua, chorionic plate, and all three combined (total) that were vRNA-positive (above the limit of detection of 3 copies/mg of tissue) at delivery. (E) PRNT90 titers one month post-infection. Box plots show the interquartile range within the box, the median as a dark horizontal line, and the minimum and maximum values excluding the outliers are shown as whiskers.

### Infant demographics and growth

A total of 41 pregnancies and their offspring were included in this study. Because there were no significant virologic differences among the maternal infection conditions, the ZIKV-exposed offspring are grouped together for subsequent analyses and compared to the control infants and then split into inoculation groups if differences were observed between control and ZIKV-exposed offspring. The control offspring were exposed to the same prenatal and postnatal stressors (maternal sedation events, infant exams, blood draws) as the ZIKV-exposed offspring. The ZIKV-exposed and control infants had similar gestational ages at inoculation, 39.8 and 40.2 days, respectively (Table S4). Half of the ZIKV-exposed infants were male (51.7%) and none of the control infants were male (t-test; p-value 0.0014) (Table S4). The proportion of male and female infants in each group was random because dams were divided into inoculation groups before fetal sex was known. Nearly all infants were delivered by Cesarean section in both the ZIKV-exposed (89%) and control (92%) groups (Table S4). There was no significant difference in proportion of infants that were housed with a dam (either biological or surrogate) compared to infants housed in the nursery then peer group reared throughout the first year of life or infant ages at hearing, eye, or developmental examinations (Table S4). ZIKV-exposed infants have similar weights at birth (within 10 days of birth) as control infants, but have a significantly greater weight gain trajectory from birth to 12 months of age (p-value 0.0317), and are heavier at 12 months of age (p-value 0.0121) (Figure 2, Table S5). There was no association between ZIKV-exposed infants’ weight at 12 months and maternal viral or immunologic parameters (Figure S1). Head circumferences were similar between ZIKV-exposed and control infants at birth and 12 months of age, and the trajectories were similar as well (Table S5).

**Fig. 2.**
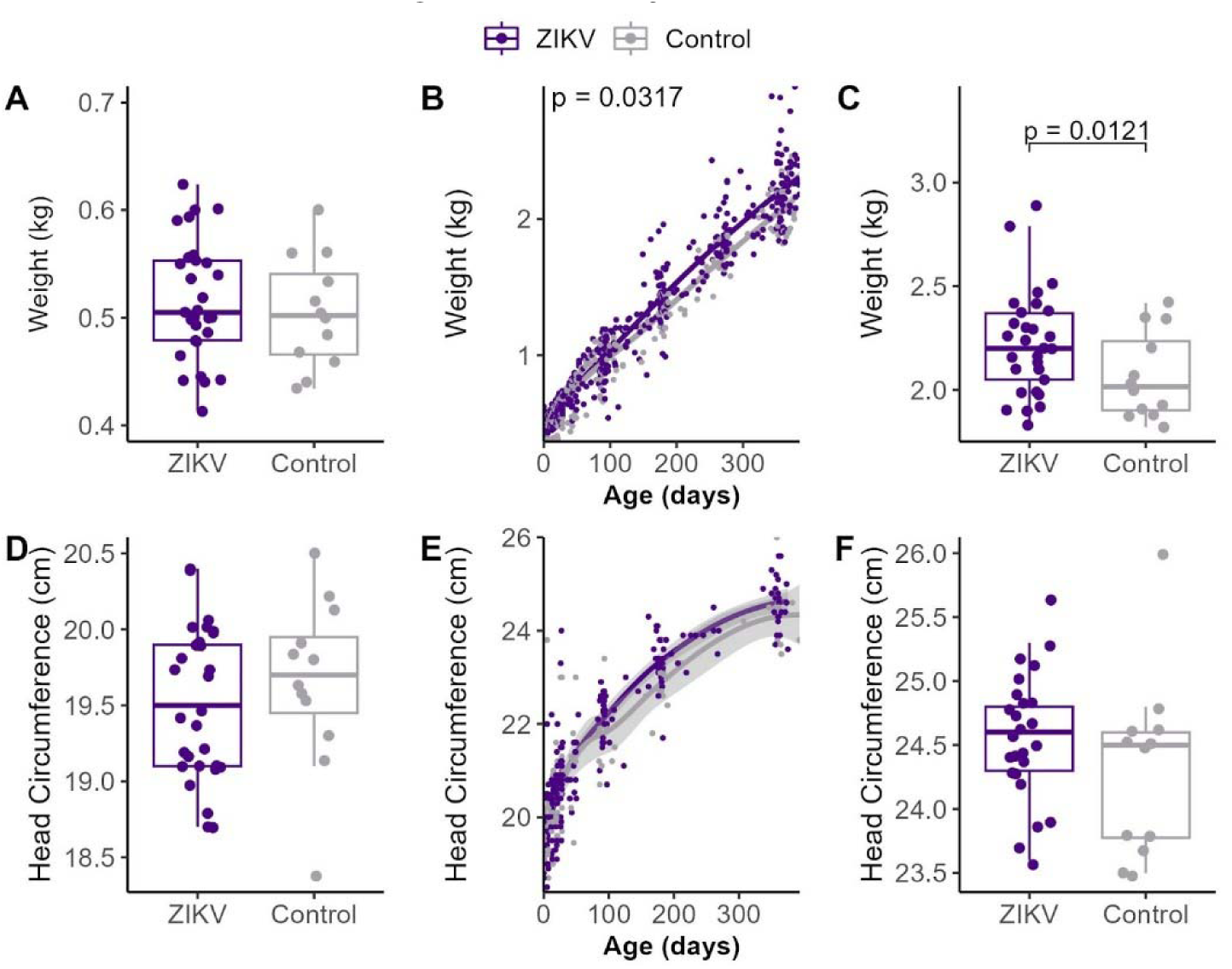
Infant growth from birth to 12 months of age. Infant weight and head circumference was measured longitudinally (ZIKV n=29, control n=12). (A) Birth weights. (B) Weight gain trajectory from birth to 12 months. The mean weight is shown by the solid line and the confidence interval by the shaded area. ZIKV-exposed infants had a significantly greater weight gain trajectory (p = 0.0317). (C) Weights at 12 months. ZIKV-exposed infants had significantly greater weights at 12 months (p = 0.0121). (D) Head circumferences at birth. (E) Head circumferences from birth to 12 months. The mean head circumference is shown by the solid line and the confidence interval by the shaded area. (F) Head circumferences at 12 months.

### Infant development

We compared infant development at 12 months of age between ZIKV-exposed infants and controls within the domains of social-emotional development, sensory responsiveness, visual motor, fine motor, and cognitive skills. Behavioral assessments included ZIKV (n=26) and control (n=9) infants raised in dyads. Due to factors such as COVID-19 research restrictions, there is missing data in some behavioral assessments. Infants raised in peer-groups were excluded from the behavioral analysis.

*<u>Mother-infant home cage observations</u>*Social-emotional development was assessed by analyzing video recordings of maternal-infant interactions in their home cage. ZIKV-exposed infants spent significantly more time in mutual ventral contact (p-value 0.0127), close proximity with their mothers (p-value 0.0149), and nipple contact (p-value 0.0093), compared to control infants (Figure 3, Table S6), suggesting that ZIKV exposure impacts social attachment. These behaviors were not mutually exclusive—infants could be in nipple contact and mutual ventral contact simultaneously—but they were not entirely co-dependent, as nipple contact sometimes occurred independently. To determine whether increased maternal contact in ZIKV-exposed infants resulted from infant impaired motor function, we compared the duration of time spent moving around the cage (i.e. in locomotion). There was no significant difference in the time spent moving through the cage between ZIKV-exposed and control groups (p-value 0.3751) (Figure 3), indicating that reduced locomotion was not responsible for increased maternal attachment. Next, we examined whether the inoculation group influenced the mutual ventral contact phenotype, a well-recognized measure of attachment from video recordings. Infants in the ZIKV-PR 45gd group spent significantly more time in mutual ventral contact than controls (p-value 0.0104), with no other significant pairwise differences (Table S6). We also assessed if maternal virologic and antibody variables predicted mutual ventral contact duration using linear regression.

**Fig. 3.**
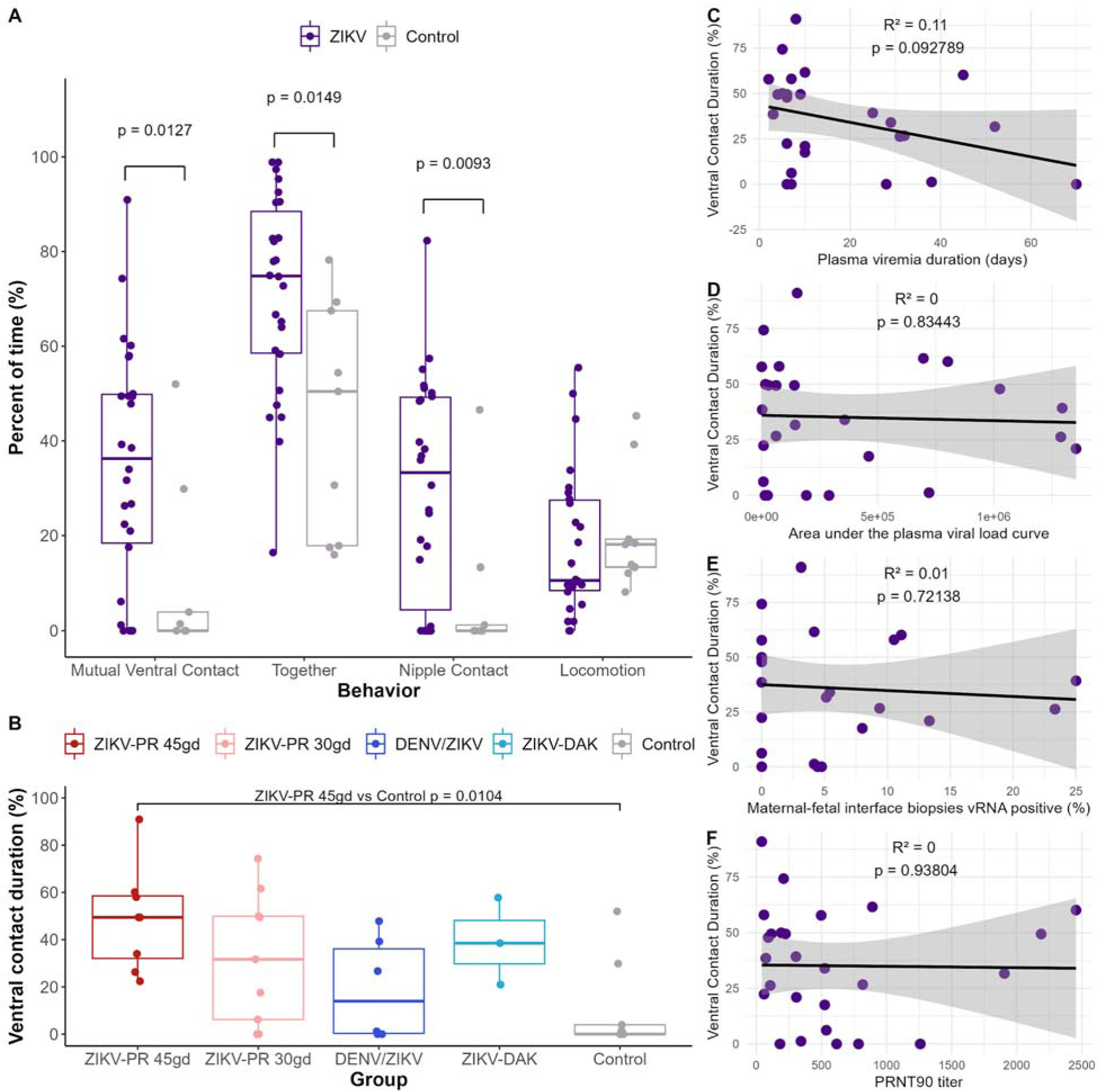
Persistent maternal-infant attachment in ZIKV-exposed infants at 12 months of age. Mother-infant dyads (ZIKV n=26, control n=9) were videotaped in their home cage, and the durations spent in nipple contact, togetherness, ventral contact, and locomotion were recorded as a percentage of the total observation time (A). Boxplots represent the median and interquartile range for each behavior, with individual data points overlaid. Significant differences between ZIKV-exposed and control groups are indicated with p-values: Mutual Ventral Contact (p = 0.0127), Togetherness (p = 0.0149), and Nipple Contact (p = 0.0093). (B) Mutual ventral contact duration by inoculation group. A significant difference between ZIKV-PR 45gd and control groups is indicated with p =0.0104. The association between mutual ventral contact duration and maternal infection variables including plasma viremia duration (C), area under the curve plasma viral load (D), maternal-fetal interface biopsies that were vRNA positive (E) and PRNT90 titer (F) were defined.

Maternal viremia duration, area under the viral load curve, percentage of maternal-fetal interface biopsies positive for vRNA, and PRNT90 titer were not significantly associated with mutual ventral contact duration (Figure 3). These findings suggest that prenatal ZIKV exposure alone influences social attachment development at 12 months of age, but specific maternal virologic and antibody measurements did not.

#### Puzzle feeder

Infant fine motor, visual motor, and cognitive skills were assessed through manipulation of the puzzle feeder over three days. The maximum puzzle feeder level completed by either group was 4 out of 8. The ZIKV-exposed infants performed similarly as the control infants for the number of levels completed (Figure 4, Table S6). Although they ended up completing a similar number of levels, some ZIKV-exposed infants took longer to complete level 1, even though there was no significant difference in the time to complete level 1 overall (Figure 4). ZIKV-exposed infants had a similar number of attempts to complete level 1 and level 2 as the control infants (Figure 4). To assess the role of motor deficits, digit isolation and overall motor coordination were assessed, but no significant differences were identified (Table S6). These findings indicate that prenatal ZIKV exposure does not impair motor, visuomotor, or cognitive performance on a puzzle feeder task at 12 months of age in this study.

**Fig. 4.**
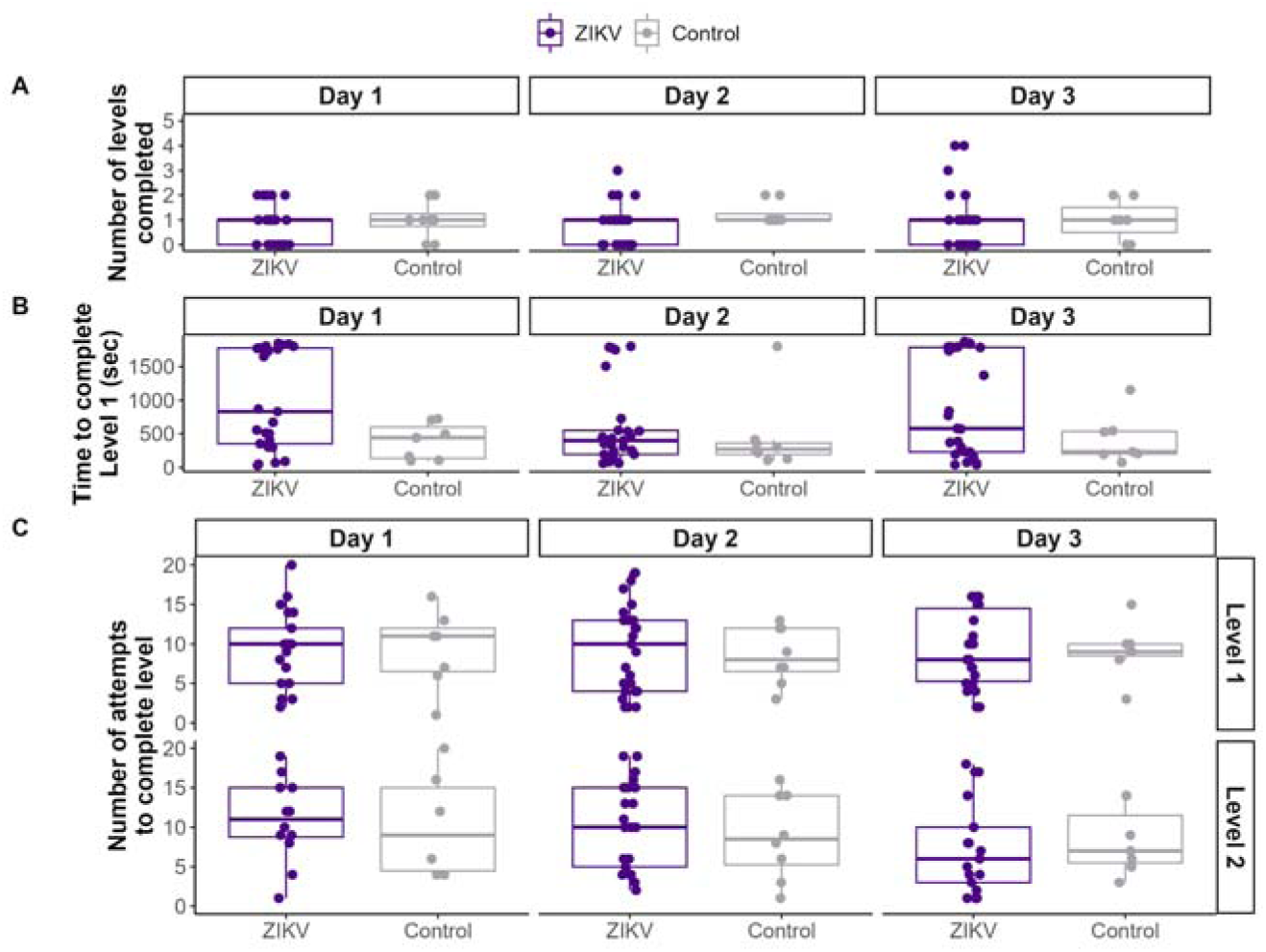
Puzzle feeder task evaluation of fine motor, visual motor, and cognitive skills. (A) The number of levels completed by ZIKV-exposed and control infants did not differ. (B) The time to complete level 1 did not differ significantly between groups, but some ZIKV-exposed infants had much higher times to complete level 1. (C) The groups displayed a similar number of attempts to complete level 1 and 2.

#### Fine Motor PVC Pipe Test

Infant fine motor skills and sensory responsiveness were assessed using the PVC Pipe Test that included manipulating a PVC pipe that was filled with raisins and frosting. The ZIKV-exposed infants did not differ in the percentage of time that they orally explored the materials or manipulated the frosting with their fingers. In addition, they did not differ on whether they were isolating their fingers to explore the pipe and raisins (Figure S2, Table S6).

#### Sensory Processing Measure for Monkeys

We also defined infant sensory processing response by assessing their approach to three sensory stimuli (1. feather, 2. cotton ball, 3. brush) in the same order over three consecutive days of testing. The duration of time that it took each infant to approach the sensory stimuli was categorized into no approach, >2 minutes, 1-2 minutes, or <1 minute. On the first day of testing, most of the ZIKV-exposed infants approached the sensory stimuli, whereas only about half of the control infants approached each stimulus (brush p-value 0.0374, cotton ball p-value 0.0035, feather p-value 0.0085). Of the ZIKV-exposed infants that approached the stimulus on day 1, most approached quickly within 1 min, with fewer infants taking 1-2 or >2 min to approach (Figure 5A). After becoming familiarized with the stimuli on days 2 and 3 of testing, ZIKV-exposed infants had an approach pattern that became more similar to the control infants, with most approaching the stimuli quickly. To explore possible contributions of maternal viral and antibody variables to the abnormal day 1 approach phenotype for ZIKV-exposed infants, we selected the feather stimulus as the representative variable because it was most different from all normal stimuli in the cage. The proportion of infants that approached the feather stimuli differed by inoculation group, with 100% of DENV/ZIKV and ZIKV-DAK infants approaching the stimuli, compared to between 75-90% of ZIKV-PR 45gd and ZIKV-PR 30gd approaching the stimuli (Figure 5B). The percent of control infants that approached the feather stimulus was <50%. Maternal viremia duration, area under the viral load curve, percentage of maternal-fetal interface biopsies positive for vRNA, and PRNT90 titer were not significantly different between ZIKV-exposed infants that did and did not approach the feather on day 1 (Figure 5C-F).

**Fig. 5.**
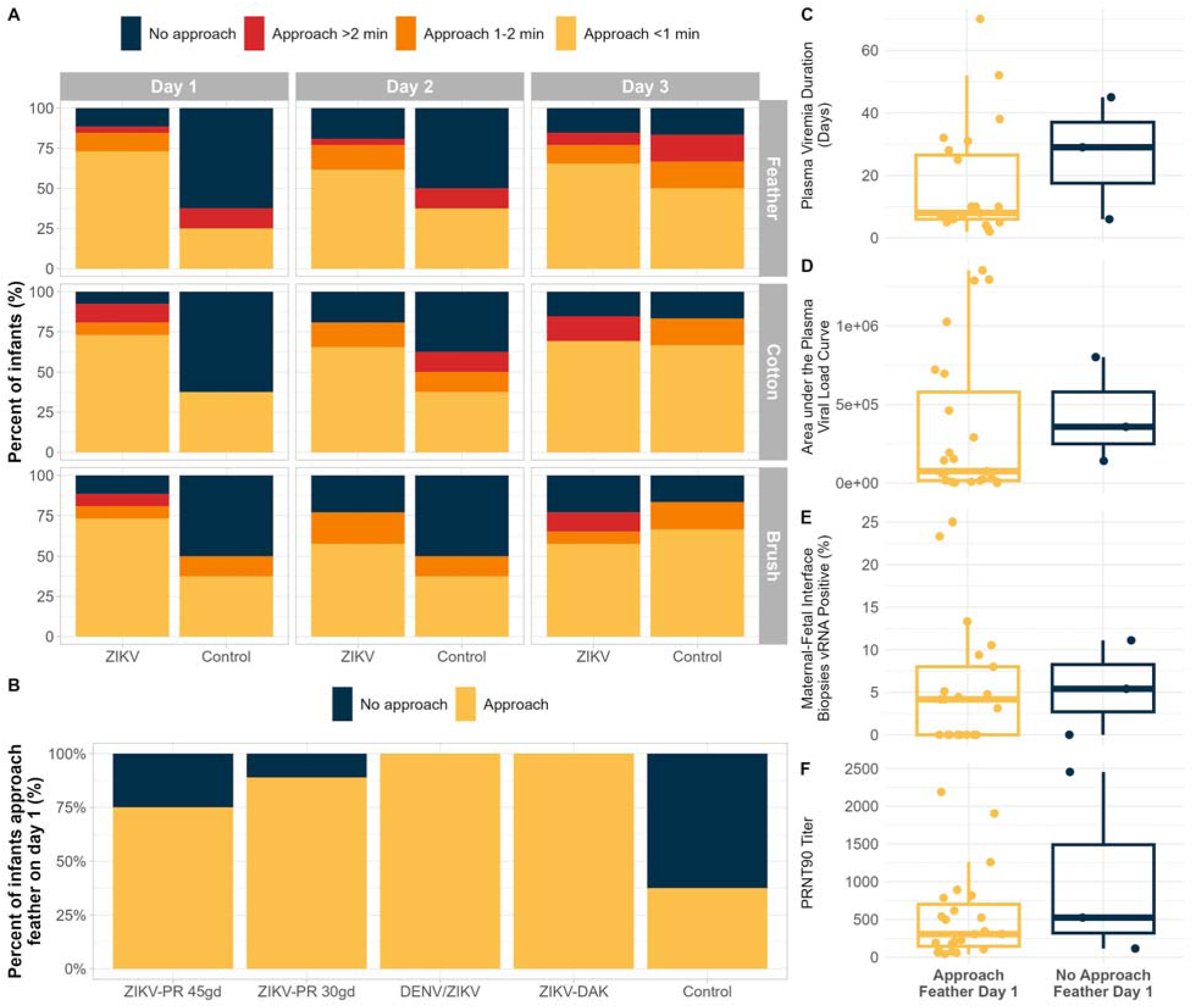
Approach to sensory stimuli differed in ZIKV-exposed and control infant macaques. Infant macaques were exposed to sensory stimuli for 3 consecutive days and their time to approach time was measured (ZIKV n=28, control n=8). (A) More ZIKV-exposed infants approached the three sensory stimuli on day 1 compared to control infants (brush p-value 0.0374, cotton ball p-value 0.0035, feather p-value 0.0085), and these differences resolved by day 2 and 3. Stimuli were presented in the same order each day (feather, cotton ball, brush) for 5 minutes, and their approach time was reported as No approach, >2 min to approach, 1-2 min to approach, and <1 min to approach. The approach to feather on day 1 was selected as a representative sensory stimulus for day 1 and used in subfigure B, C, D, E, F, to show how the binary response of approach or no approach did not differ based on inoculation group (B), plasma viremia duration (C), area under the curve plasma viral load (D), maternal-fetal interface biopsies that were vRNA positive (E) and PRNT90 titer (F).

Altogether, this indicates that ZIKV-exposed infants initially had decreased inhibition when approaching sensory stimuli in their cage.

### Infant vision and hearing development

*<u>Structural eye evaluation</u>* In a prior study using this macaque model of ZIKV infection, we identified structural ocular abnormalities in a ZIKV-exposed fetus (*45*). In the present cohort of prenatally ZIKV-exposed infants, we did not observe structural eye abnormalities classically associated with congenital ZIKV infection during ophthalmic examinations. However, we identified two ocular findings not previously reported in human infants with congenital ZIKV infection—iris nodules and persistent fetal vasculature (Figure S3, Table S7). These findings may be incidental and are not expected to impact visual function. Retinal structure was assessed by measuring layer thickness using spectral-domain optical coherence tomography. Total retinal layer thickness at the center of the fovea did not differ between ZIKV-exposed and control infants. Of the eight retinal layers measured, only the outer plexiform layer showed a small, non-significant difference (Figure S4, Table S8). Together, these findings indicate that these prenatally ZIKV-exposed infants did not have structural ocular abnormalities detectable by ophthalmic examination or optical coherence tomography.

*<u>Visual Electrophysiology</u>* We evaluated retinal and cortical visual function using visual electrophysiology studies. Retinal function was assessed by electroretinography (ERG), and cortical visual pathway function was assessed by visual evoked potentials (VEPs). Retinal visual function, as measured by ERG, was comparable between ZIKV-exposed and control infants at both 3 and 12 months of age (Figure 6). In contrast, VEP responses in ZIKV-exposed infants differed significantly from controls. At 3 months of age, ZIKV-exposed infants exhibited significantly lower VEP amplitudes, as measured by the root mean square amplitude across left and right lateralities (Figure 6). These differences resolved by 12 months of age, with VEP amplitudes no longer significantly different from controls (Figure 6), suggesting delayed maturation of the cortical visual pathway in ZIKV-exposed infants. No significant associations were identified between 3-month VEP amplitudes and maternal viro-immunologic parameters, including viremia duration, area under the viral load curve, percentage of maternal-fetal interface biopsies positive for vRNA, or PRNT90 titer (Figure S5).

**Fig. 6.**
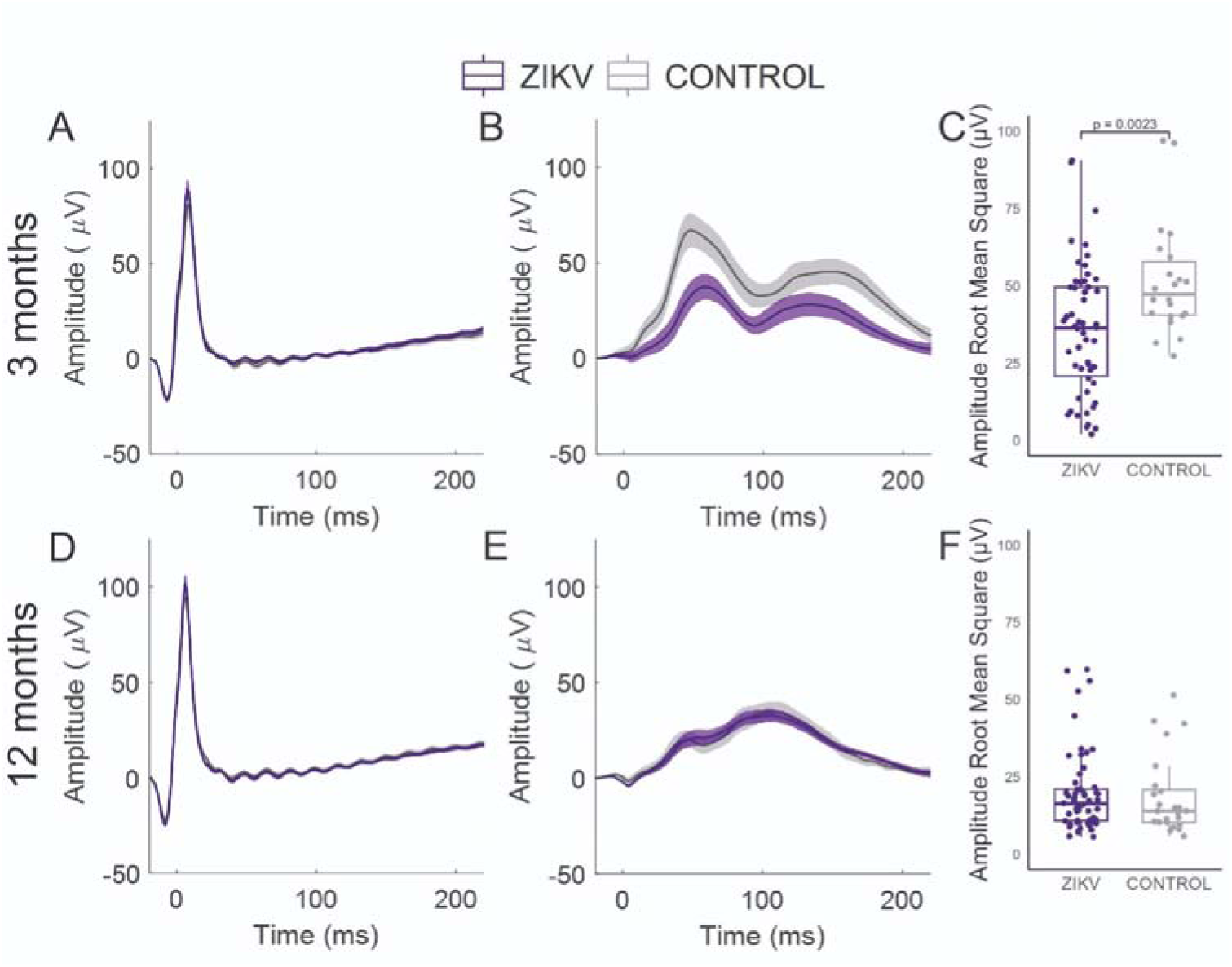
Visual electrophysiology studies for ZIKV-exposed and control infants. (A) Light-Adapted Electroretinogram (LA ERG) at 3 months. Waveforms shown are group averages (±SEM) of the left eye ERG recorded from a corneal electrode to monocularly-delivered 2.5 cd·s·m^-2^ white flashes on a 30 cd·m^-2^ background following 10 min of light adaptation. (B) Flash visual evoked potential (FVEP) group averages (±SEM) at 3 months. FVEPs were recorded concurrently with the LA ERG. Waveforms shown are group averages of combined right and left occipital FVEPs following left eye stimulation. LA ERG and FVEP waveforms elicited by right eye stimulation are not shown as they were indistinguishable from those of the left eye. (C) Root mean square (RMS) amplitudes of the FVEP from left and right occipital sites. RMS amplitudes differed between groups at 3 months of age (Wilcoxon rank-sum test, p = 0.0023). (D) LA ERG group average waveforms (±SEM) at 12 months. (E) FVEP group average (±SEM) waveforms at 12 months. (F) FVEP RMS amplitudes for ZIKV-exposed became statistically indistinguishable from controls at 12 months (p = 0.52).

*<u>Hearing</u>* We observed a higher rate of hearing loss in ZIKV-exposed infants compared to control infants at 12 months of age, although this difference did not reach statistical significance. At 12 months, 5 of 29 (17.2%) ZIKV-exposed infants and 1 of 12 (8.3%) control infants demonstrated hearing loss (Figure S6). Hearing loss was identified in the ZIKV-PR 45gd, ZIKV-PR 30gd, and DENV/ZIKV groups, but not in the ZIKV-DAK group, possibly due to the smaller sample size. Hearing loss was frequency specific, occurring with either the low pitch (i.e. low-frequency 500 Hz) or mid pitch (i.e. mid-frequency 1000 Hz) tone bursts, but not with the click stimulus, which assesses a broader frequency spectrum (Table S9). Auditory brainstem responses were absent at the lowest volume assessed (40 decibels) and not the higher volumes, suggesting mild to moderate loss. To evaluate whether the hearing loss were persistent, suggesting early-onset sensorineural hearing loss rather than transient conductive or late-onset sensorineural loss, we examined auditory brainstem responses at earlier time points for infants with hearing loss at 12 months. None of these affected infants had hearing loss immediately preceding evaluation; however, some had losses detected at earlier ages, precluding clear differentiation between transient conductive loss and late-onset sensorineural hearing loss (Table S9).

### Maternal Predictors and Potential Mediators of ZIKV-Associated Developmental Deficits

We evaluated whether developmental deficits observed in ZIKV-exposed infants were mediated vision impairments or hearing loss. Specifically, we assessed whether maternal infection characteristics—ZIKV exposure status, duration of plasma viremia, or neutralizing antibody titer—predicted two behavioral outcomes: mutual ventral contact duration and approach behavior to a novel stimulus. We selected two candidate mediators based on domains with significant group differences: visual function (visual evoked potential amplitude at 3 months) and hearing (hearing loss at 12 months). Because we have a relatively small sample size for this type of analysis, we used a p-value of 0.1 to define significance. We could not evaluate the impact of ZIKV inoculation groups on the mediators and outcomes because of the much smaller sample size within each inoculation group.

Both ZIKV exposure (ZIKV vs control) and the candidate mediators (vision and hearing) were individually associated with the behavioral outcomes (Table 1). These associations remained when adjusting for one another: vision and hearing predicted behavioral outcomes while controlling for ZIKV exposure, and ZIKV exposure predicted behavioral outcomes while controlling for vision and hearing. However, formal mediation analyses (Sobel test) did not identify statistically significant indirect effects of ZIKV exposure on behavior through either visual or auditory pathways). These findings suggest that visual and auditory abnormalities may co-occur with abnormal behavior but do not mediate the relationship between prenatal ZIKV exposure and the observed deficits in social-emotional development and sensory approach behavior.

**Table 1.**
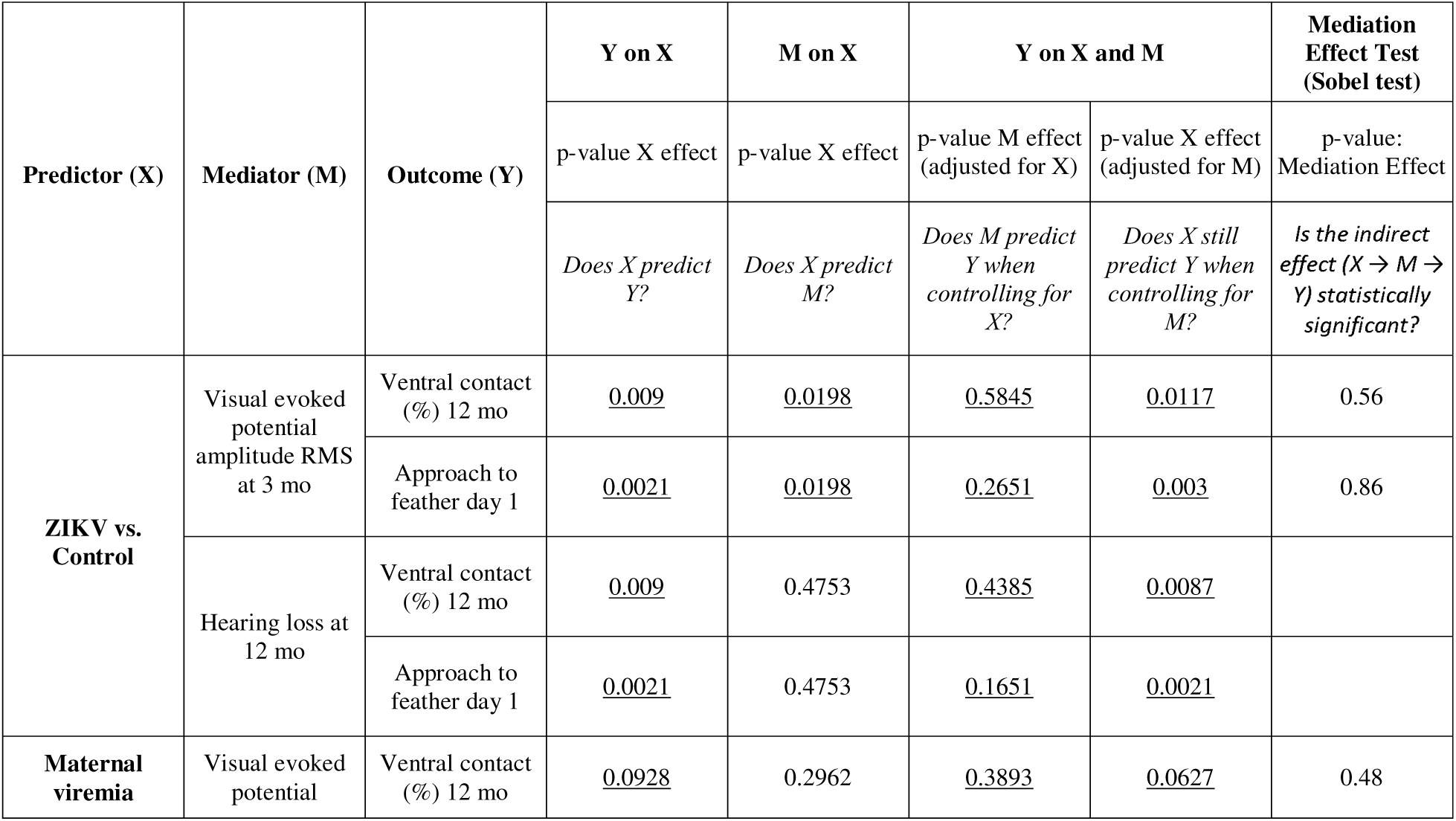
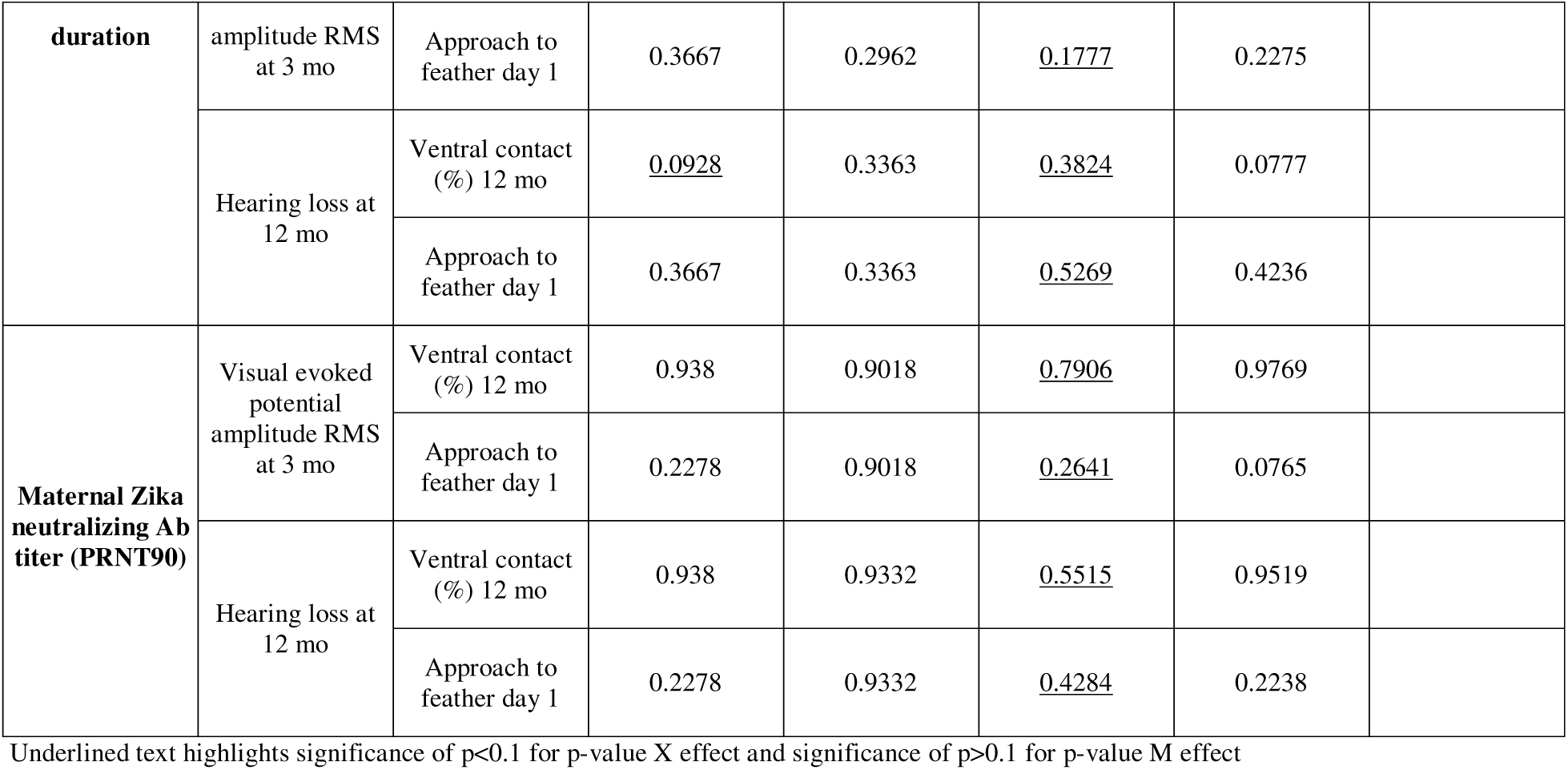
Mediation analysis

We next evaluated whether developmental deficits were mediated by vision or hearing in relation to two maternal virologic and immunologic predictors: plasma viremia duration and neutralizing antibody titer. We concentrated on these variables rather than AUC plasma viral loads and the distribution of vRNA within the maternal-fetal interface because these variables are not feasible to collect in a human population. As in the prior analysis, we focused on two developmental outcomes—mutual ventral contact duration and approach behavior—and two candidate mediators: visual evoked potential amplitude at 3 months (vision) and hearing impairment at 12 months (hearing). In these models, longer maternal viremia duration was directly associated with increased ventral contact duration but not with approach behavior, while higher neutralizing antibody titers were not significantly associated with either behavioral outcome. Although some of the mediators were associated with the outcomes, neither visual function or hearing status met criteria for statistically significant mediation between maternal viremia duration or antibody levels and the behavioral outcomes. This means that while prolonged maternal viremia may be linked to certain developmental deficits, the effect does not appear to operate through visual or, auditory differences as intermediary mechanisms.

## DISCUSSION

We aimed to define how maternal ZIKV inoculation, viremia, and antibody responses influence infant developmental trajectories. Prenatal ZIKV exposure—regardless of maternal inoculation group—increased the risk of developmental deficits, delayed maturation of the cortical visual pathway, and hearing loss. At 12 months of age, ZIKV-exposed infant macaques exhibited abnormal development across multiple domains, including social-emotional delays and reduced inhibition to sensory stimuli, compared to procedure-matched controls. This constellation of findings mirrors the mild-to-moderate neurodevelopmental impairments most commonly reported in children with prenatal ZIKV exposure (*3–8*). Importantly, these impairments were attributable to ZIKV exposure itself and were not mediated by maternal virologic or immunologic status, viral lineage, gestational timing of infection, or prior DENV immunity.

Mediation analyses further supported a direct effect of ZIKV exposure on neurodevelopment, particularly in social attachment behaviors, independent of changes in visual or auditory function. These findings demonstrate that even in the absence of overt congenital anomalies, prenatal ZIKV exposure can disrupt early neurobehavioral development. They underscore the need for risk assessment protocols that account for subclinical outcomes and support early developmental screening and longitudinal neurodevelopmental monitoring in exposed infants.

At 12 months of age, ZIKV-exposed infant macaques exhibited significant differences in social-emotional behavior, while their cognitive, fine motor, and gross motor development remained comparable to controls. ZIKV-exposed infants spent more time in mutual ventral contact, close proximity, and nipple contact with their mothers—behaviors that typically decline as infants transition toward greater independence and peer-group engagement. This increased maternal attachment may reflect underlying disruptions in emotional regulation, threat perception, or social motivation, and could interfere with the development of normative peer-directed social behaviors. Notably, the increased duration of nipple contact may also contribute to the accelerated growth trajectory observed in ZIKV-exposed infants. While similar mother-infant interactions have not previously been characterized in macaque models of congenital viral infections, altered social behavior has been described in prenatal immune activation models (*46*). Additionally, emotional reactivity is disrupted in infant macaques infected postnatally with ZIKV (*47*), suggesting that social-emotional alterations may be a shared outcome of both prenatal and postnatal ZIKV exposure. Together, these findings indicate that prenatal ZIKV exposure disrupts the normative trajectory of social-emotional development, increasing maternal dependence and potentially hindering peer engagement.

ZIKV-exposed infants also demonstrated a lack of inhibition, characterized by an accelerated approach to sensory stimuli introduced into their environment. This pattern may reflect inappropriate approach behaviors or impaired sensory processing and integration, leading to deficits in novelty detection or threat assessment. Similar exaggerated or unmodulated responses to environmental stimuli have been reported in macaques exposed to prenatal stress (*48*), prenatal lead (*49*), and maternal ZIKV infection (*50*). These neurobehavioral features may serve as early indicators of neurodevelopmental risk for later-emerging deficits in social behavior, anxiety regulation, or cognitive flexibility (*51*). A lack of inhibition may also suggest delayed or atypical emotional learning, resulting in diminished caution or blunted responsiveness to adverse cues. Collectively, these findings implicate potential disruption of brain regions critical to inhibitory control and emotional regulation, such as the prefrontal cortex and amygdala. They further raise the possibility that behavioral phenotypes observed in ZIKV-exposed children— such as altered attachment or sensory processing disorders—may have neurobiological origins detectable in early infancy.

Human studies have shown that prolonged maternal viremia (>30 days) is associated with increased risk of fetal loss and structural brain abnormalities (*24*), while higher maternal neutralizing antibody titers two months post-infection are linked to reduced risk of microcephaly and structural brain abnormalities (*28*). In our macaque cohort, consistent with prior studies [reviewed in (*20*)], we did not observe microcephaly, limiting our ability to assess whether maternal viral or immunologic parameters are associated with these more severe outcomes— often considered the "tip of the iceberg" in congenital ZIKV infection. Notably, we found that prenatal ZIKV exposure increased the risk of developmental deficits and cortical visual pathway delays regardless of maternal viremia duration or antibody responses. This challenges the prevailing assumption that longer maternal viremia necessarily leads to worse neurodevelopmental outcomes. Despite a wide range of viremia durations (2–70 days) in our cohort, no association was observed between viremia length and infant outcomes. These findings highlight the need for prospective human studies to determine whether longer maternal viremia is indeed predictive of later developmental deficits and, in turn, whether antiviral treatment can improve long-term neurodevelopment. Ultimately, prevention strategies—such as immunization or mosquito control—may prove more effective for improving outcomes than treatment after infection.

ZIKV-exposed infants in our study showed delayed maturation of the cortical visual pathway despite normal ocular structure and retinal signaling. While retinal abnormalities are a hallmark of congenital Zika syndrome, less is known about visual function in infants without structural eye defects. We found significantly reduced visual evoked potential (VEP) amplitudes in ZIKV-exposed infants, suggesting dysfunction in the post-retinal visual pathway to the occipital cortex. In contrast, electroretinography and retinal imaging were largely normal, and rare findings such as iris nodules and persistent fetal vasculature were unlikely to affect vision. Similar patterns have been observed in children with congenital ZIKV infection, where VEP abnormalities occur despite normal ophthalmic exams (*52*), or alongside retinal changes with primarily latency shifts (*53*). Cortical visual impairment, a leading cause of childhood visual dysfunction, can affect acuity and higher-order processes such as motion perception and face recognition (*54*). Although early cortical visual deficits were detected at 3 months in our cohort, they resolved by 12 months, and we found no association between early visual function and later developmental outcomes. Ongoing analyses will examine how changes in cortical visual function across the first year of life relate to neurodevelopmental trajectories from birth to 12 months. These findings underscore the importance of comprehensive visual assessments in ZIKV-exposed children, even in the absence of structural abnormalities, and point to the need for further study of subtle or transient cortical visual impairments.

This study has several important limitations. First, although the sample size is large for a nonhuman primate model, it may have limited our power to detect subtle group differences or mediation effects, particularly for binary outcomes such as hearing loss. This is especially relevant given that approximately 30% of infants are expected to exhibit neurodevelopmental deficits, consistent with rates reported in human cohorts (*3*, *4*). The limited sample size within each inoculation group also precluded mediation analyses examining how factors such as timing of maternal ZIKV exposure, prior DENV immunity, or viral lineage influence developmental outcomes through intermediate variables. Second, although we evaluated a broad range of developmental domains, some behavioral or cognitive impairments may emerge later in life and would not have been detected at the 12-month time point. Despite these limitations, this remains the largest study of nonhuman primate development following prenatal ZIKV exposure (*20*), and the only one to date to examine associations between maternal virologic and immunologic characteristics and infant developmental outcomes.

In summary, our findings demonstrate that prenatal ZIKV exposure results in measurable neurodevelopmental deficits—including altered social-emotional behavior, sensory processing abnormalities, hearing loss, and delayed cortical visual pathway maturation—even in the absence of structural birth defects. These outcomes were not predicted by maternal viremia duration, antibody titers, ZIKV lineage, or DENV immunity, challenging the assumption that maternal virologic or immunologic features alone can accurately stratify fetal risk. This highlights a critical limitation in current maternal-centric approaches to evaluating the effectiveness of prenatal interventions. Infant-focused outcome measures—particularly neurodevelopmental endpoints—must be incorporated into the design and evaluation of maternal treatment trials to capture the full spectrum of congenital ZIKV impact. The combination of behavioral, electrophysiologic, and structural assessments used in this translational model provides a framework for identifying early markers of atypical development and evaluating the downstream effects of maternal therapies. These findings also support the need to include all ZIKV-exposed infants—regardless of symptom status at birth—in pediatric developmental screening and follow-up, to enable early identification of delays and timely intervention during critical periods of early childhood development.

## MATERIALS AND METHODS

### Study design, inoculation, and monitoring

Indian-origin rhesus macaques (Macaca mulatta) were inoculated with ZIKV or phosphate-buffered saline (PBS) during the first trimester, either early in the first trimester around 30 gestational days (gd) or closer to the second trimester around 45gd. All dams were housed at the Wisconsin National Primate Research Center (WNPRC) and were free of Macacine herpesvirus 1 (Herpes B virus), simian retrovirus type D (SRV), simian T-lymphotropic virus type 1 (STLV), and simian immunodeficiency virus (SIV). To generate pre-existing immunity to DENV, eight macaques were inoculated with 1x10^4^ plaque forming units (PFU) DENV-2/US/BID-V594/2006 37-68 days prior to breeding, as previously described (*55*). Macaques were bred, and once pregnant, were inoculated subcutaneously over the cranial dorsum with PBS, or 1x10^4^ PFU Zika virus/H.sapiens-tc/PUR/2015/PRVABC59_v3c2 (PRVABC59, GenBank: KU501215), or 1x10^8^ PFU Zika virus/A.africanus-tc/Senegal/1984/DAKAR 41524 (ZIKV-DAK; GenBank: KX601166) (Table S10). Preparation of the ZIKV-PR (*23*) and ZIKV-DAK (*42*) virus stocks have been described previously. The pregnancies of the seven DENV/ZIKV and four ZIKV-DAK pregnancies have been described earlier (*43*, *55*). Post-inoculation, the animals were closely monitored by veterinary and animal care staff for adverse reactions or any signs of disease. Blood was drawn for ZIKV RT-qPCR (or neutralizing antibody titers) daily for 10 days following inoculation during pregnancy, then twice weekly until viremia cleared, then weekly until Cesarean delivery. Infants had blood drawn for ZIKV RT-qPCR immediately after delivery, or within the first week of life, if the infant was born naturally, and with other sedated exams (overview in Figure 7; details in Table S10).

**Fig. 7.**
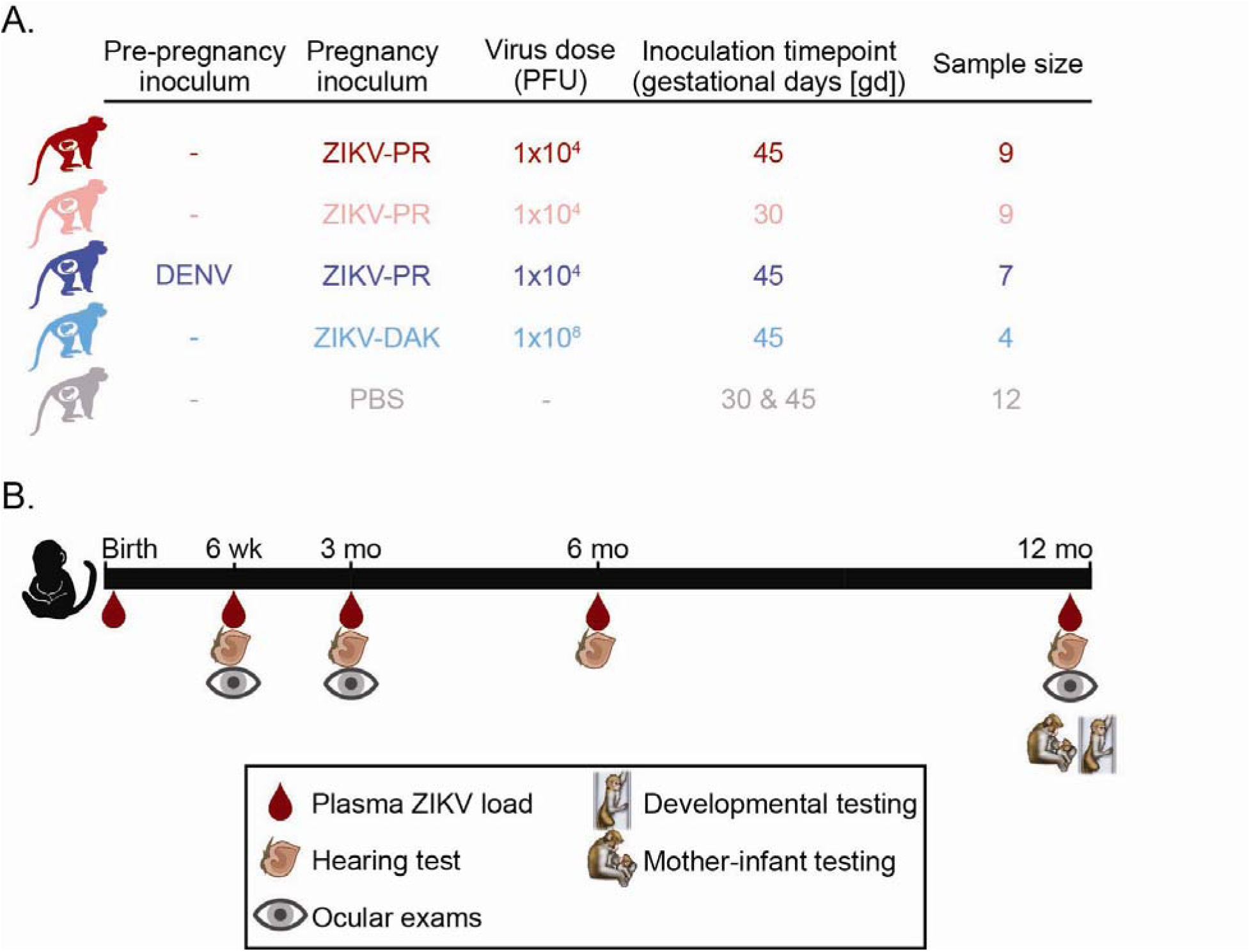
Experimental timeline schema. (A) Pregnant macaques were inoculated with ZIKV-PR, ZIKV-DAK or PBS at approximately 30 or 45 gestational days. Some females had a history of DENV infection 1-2 months prior to pregnancy. Specific inoculation virus and timeline are described in Table S10. Inoculation group colors are consistent throughout the manuscript. (B) After delivery, infants underwent a battery of tests, including ZIKV plasma viral loads, IgM testing, hearing tests, ocular exams, developmental testing and mother-infant observations at the time points indicated. Created with Biorender.com.

### Cesarean delivery

Infants were delivered by Cesarean at approximately 160 gestational days (gd) (+/- 5 days) (full details in Table S11), about 8 days earlier than the average gestational age of a natural birth (*56*) to ensure that the placenta could be collected for virologic and histologic evaluation. Full thickness sections of each cotyledon in each placental disc were collected for vRNA isolation from the villous parenchyma, decidua basalis, and chorionic membranes. Amniotic fluid was collected immediately prior to infant delivery using a syringe and needle inserted through the fetal membranes. Two infants were delivered through spontaneous parturition (044–108, 044–109, 044–531) prior to their scheduled Cesarean section; placental tissues could not be collected from 044-109.

### Ethics statement

All rhesus macaques are cared for by the staff at the WNPRC in accordance with the regulations and guidelines outlined in the Animal Welfare Act and the Guide for the Care and Use of Laboratory Animals, the recommendations of the Weatherall report (*57*), and the principles described in the National Research Council’s Guide for the Care and Use of Laboratory Animals (*58*). The University of Wisconsin - Madison Institutional Biosafety Committee approved this work under protocol number B00000764. See study approval section below for animal protocol details.

### Care & use of macaques

All animals were housed in enclosures with required floor space and fed using a nutritional plan based on recommendations published by the National Research Council (*58*). Dams were fed a fixed formula, extruded dry diet with adequate carbohydrate, energy, fat, fiber, mineral, protein, and vitamin content. Macaque dry diets were supplemented with fruits, vegetables, and other edible objects (e.g., nuts, cereals, seed mixtures, yogurt, peanut butter, popcorn, marshmallows, etc.) to provide variety to the diet and to inspire species-specific behaviors such as foraging.

When needed, infants were fed 5% dextrose for the first 24 hours of life and liquid formula subsequently. To further promote psychological well-being, animals were provided with food enrichment, structural enrichment, and/or manipulanda. Environmental enrichment objects were selected to minimize chances of pathogen transmission from one animal to another and from animals to care staff. While on study, all animals were evaluated by trained animal care technicians at least twice each day for signs of pain, distress, and illness by observing appetite, stool quality, activity level, and physical condition. Animals exhibiting abnormal presentation for any of these clinical parameters were provided appropriate care by attending veterinarians. Prior to all minor/brief experimental procedures, macaques were sedated using ketamine anesthesia and monitored regularly until fully recovered from anesthesia.

For breeding, the female macaques were co-housed with a compatible male and observed daily for menses and breeding. Pregnancy was detected by ultrasound examination of the uterus at approximately 20-24 gestation days (gd) following the predicted day of ovulation. The gd was estimated (+/- 2 days) based on the dam’s menstrual cycle, observation of copulation, and the greatest length of the fetus at initial ultrasound examination which was compared to normative growth data in this species (*59*). For physical examinations, virus inoculations, ultrasound examinations, blood or swab collections, the dam was anesthetized with an intramuscular dose of ketamine (10 mg/kg). Blood samples from the femoral or saphenous vein were obtained using a vacutainer system or needle and syringe. Pregnant macaques were monitored daily prior to and after viral inoculation for any clinical signs of infection (e.g., diarrhea, inappetence, inactivity, fever and/or atypical behaviors).

### vRNA isolation from blood and tissues and RT-qPCR

RNA was extracted from 300 µL of plasma using the Viral Total Nucleic Acid Purification kit (Promega, Madison, WI, USA) on a Maxwell 48 RSC instrument. RT-qPCR was performed as previously described (*60*). Fetal and maternal–fetal interface tissues (placenta, decidua, umbilical cord, chorionic plate, fetal membranes) were preserved with RNAlater® (Invitrogen, Carlsbad, CA, USA) at 4°C for 24–72 h before the RNAlater was removed and the tissue was frozen at −80°C. RNA was isolated from maternal and fetal tissues using a method described by Hansen et al. (*61*) and previously described in detail (*22*). The RT-qPCR limits of detection are 150 copies/mL from plasma and estimated to be 3 copies/mg from tissue. The duration of maternal plasma viremia was defined as the last RNA viral load equal to or greater than the RT-qPCR limit of detection of 150 copies/mL. The distribution of vRNA within the maternal-fetal interface was defined as the fraction of ZIKV vRNA-positive biopsies divided by the number of biopsies assessed for viral loads within the placenta, decidua and chorionic plate.

### Plaque Reduction Neutralization Test (PRNT)

Neutralizing antibody titers were measured in serum samples one month post-infection (range 24-38 days) using a plaque reduction neutralization test using a 90% cutoff (PRNT90). Endpoint titrations of reactive sera were performed against ZIKV-PR for dams inoculated with ZIKV-PR (ZIKV-PR 30gd, ZIKV-PR 45 gd, DENV/ZIKV), or against ZIKV-DAK for dams inoculated with ZIKV-DAK, as previously described (*44*, *55*, *62*). These data were previously published for the ZIKV-PR 30gd and 45gd dams (*62*), for the DENV/ZIKV dams (*55*), and for the ZIKV-DAK dams (*44*).

### Infant care

After delivery, infants were dried, stimulated, and received respiratory support as clinically indicated. Birth weight was measured shortly after delivery. Infants were returned to their biological dam following her recovery from anesthesia. If the dam rejected the infant after multiple reintroduction attempts over the subsequent 2–3 days, the infant was placed with a surrogate dam from the colony when available or transferred to the nursery if no surrogate was available. Upon reaching 1 month of age, infants (n=6) initially raised in the nursery were placed in an age-matched peer group and remained with this peer group until 12 months of age. Infants in peer-raised groups were not included in behavioral analysis. Infants (n=35) initially placed with a dam remained with the same dam until 12 months of age. Infant weights were measured when dam and infant were separated for any reason. Head circumference was measured at each infant assessment using a head circumference tape measure. When multiple measurements (up to three) were obtained, the median value was recorded.

### Mother-infant home cage observations

Mother-infant dyad behavior was assessed using a 25 to 30-minute home-cage video recording at 12 months of age to evaluate the infant’s social-emotional, motor, feeding, self-care, and exploratory behaviors in a familiar environment. An observer known to the animals was present when recording the dyad to adjust the camera as needed. Behaviors were analyzed for two consistent time segments of five minutes each from near the beginning and end of the observation to capture the range of behaviors. Infants in peer groups were not included in this assessment as mother-infant dyad interactions could not be measured; this resulted in the exclusion of three ZIKV-exposed and three control infants. Mutual ventral contact, nipple contact, proximity to mother (together), and infant locomotion were coded (Table S12).

Behaviors were not mutually exclusive and could be coded simultaneously as appropriate.

### Puzzle feeder

Infants’ fine motor, visual motor, and cognitive skills were assessed through the macaques’ manipulation of a treat through a 9-level puzzle in their home cage at 12 months of age. The puzzle feeder, a clear Plexiglas puzzle that has 9 levels of increasing complexity, was attached to the side or front of the home cage enclosure (Figure S7). A food treat was placed in the top compartment and the infant must move the treat laterally through multiple paths to reach the bottom of the puzzle feeder, where the treat can be removed from a larger opening in the feeder. Two puzzle feeders were attached to opposite sides of the cage, so that the mother and infant could be engaged in separate puzzle feeders simultaneously. The infant’s interaction with the puzzle feeder was recorded by a tester familiar to the dyad using two stationary cameras, each providing a full view of one of two puzzle feeders attached to the sides of the dyad’s home cage. The mother and infant had 30 minutes to engage in the task. Both puzzle feeders were set at the easiest level first and then advanced to more difficult levels as the previous level was successfully completed. Completion of levels, individual attempts, digit isolation, and motor coordination were coded (Table S12).

### Fine Motor PVC Pipe Test

Infants were observed in their home cage to define fine motor skills and sensory differences using a 10 minute PVC pipe enrichment test at 12 months of age. The observation was recorded by a familiar tester, using both a handheld camera and a stationary camera. Three PVC pipes, approximately 0.5 inches in diameter and 4 inches in length, were prepared by filling both ends with frosting and layering in three raisins at each end to provoke the use of fine motor movements. One PVC was completely inserted into the cage to engage the mother while the second PVC was inserted into the cage frame, stabilized between the bars to more easily capture infant interactions. If the mother monopolized both PVCs, a third PVC was held next to the cage frame by the tester to capture infant interactions. The assessment was recorded for 10 minutes on three sequential days; the first day of testing was considered acclimation to task and wasn’t analyzed except to substitute for missing data on a subsequent test day for (n=2, 1 ZIKV, 1 control) infants. Oral exploration, frosting exploration, and digit isolation were coded (Table S12). All pipes remained in the cage a minimum of 98% of the time during the task.

### Sensory Processing Measure for Monkeys

Infant responses to sensory materials were assessed in their home cage environment at 12 months of age using the Sensory Processing Measure for Monkeys (*48*). Three sensory stimuli, feathers (black colored), cotton balls, or a brush, were attached to a metal rod and presented sequentially in the cage independently and in the same order each time, for a duration of five minutes per stimulus for a total of 15 minutes. Exposure to sensory stimuli was shortened if the infant or dam demonstrated excessive aggression towards stimuli such as destroying or eating materials. Infant approach, time to approach, and method of approach (visual or touch) were coded (Table S12).

### Coding reliability and software

A video coding software program, Noldus Observer XT 14.0 was used to code infant behaviors for the mother-infant dyad interactions and Fine Motor PVC Pipe test. Qualtrics (Qualtrics, Provo, UT) was used to capture the coding for the Puzzle Feeder and Sensory Response tests. All videos for all behavior protocols were coded by two individuals and an inter-rater reliability of Cohen’s kappa=0.9 was used. Any behaviors with a kappa below 0.9 were reviewed by the specific coding team until consensus was reached. All coders were blinded to the infant’s group.

### Auditory brainstem response testing

Hearing, or auditory brainstem function, was assessed with auditory brainstem response audiometry, which measures brainstem evoked potentials generated by a brief click, as previously described (*22*). Examinations were performed by a human audiologist with pediatric training (A. Hartman) who was blinded to the infant’s inoculation group. Auditory brainstem response (ABR) thresholds were obtained for auditory (click) stimuli and tone bursts (500 and 1000 Hz), as described in (*63*), using the Biologic Navigator Pro system. Ambient noise level was minimized. Needle electrodes were placed at the brow ridge (positive input) and behind the right pinna (negative input) for channel 1 and from the brow ridge (positive input) and behind the left pinna (negative input) for channel 2. An electrode was placed below the brow ridge on the forehead for the ground. Electrode impedances were below 10 kohm for all electrodes.

Physiological filters were set to pass 100–3000 Hz. The stimuli were clicks with rarefaction, condensation, and alternating polarities and tone bursts with alternating polarity gated with a Blackman window with 2 ms rise-fall and 1 ms plateau times. Insert earphones (Etymotic ER- 3A) were used for testing. Signal levels for click stimuli were presented at 70, 50, 30, and 20 dB nHL and wave IV was observed at each presentation level. Signal levels for tone burst stimuli were presented at 80, 60, and 40 dB nHL and wave IV was observed at each presentation level. Each response was recorded twice for reliability, with the noise floor being less than 10% of sweeps. To distinguish sensorineural hearing loss, which is typically persistent, from conductive hearing loss, which is often transient, infants with absent wave IV responses to one or more stimuli at 12 months of age were identified. Longitudinal wave IV responses from 1 to 12 months were then reviewed for these infants to assess for persistent patterns of hearing deficits.

### Ophthalmic exam

Ophthalmic exams were performed by a human ophthalmologist with retinal fellowship training (M. Nork) who was blinded to the infant’s inoculation group. Slit-lamp biomicroscopy and indirect ophthalmoscopy were performed after pupillary dilation.

### Optical coherence tomography

Spectral-domain optical coherence tomography is a noninvasive imaging method that uses reflected light to create images of the retinal layers. High resolution scans were collected through the center of the fovea and around the optic nerve. Scans of the retina were carried out in both eyes using a Heidelberg™ Spectralis HRA + OCT (Heidelberg™ Engineering, Heidelberg, Germany) instrument. Determination of retinal layer thickness was performed using combined manual and automatic segmentation algorithms from both Heidelberg and EXCELSIOR Preclinical (EdgeSelect™)(*64*). After segmentation, EXCELSIOR Preclinical was used to calculate mean thicknesses for individual retinal layers.

### Visual electrophysiology

To objectively evaluate visual function, ocular specialists under the auspices of a clinically trained visual electrophysiologist (J. Ver Hoeve) performed standard visual electrodiagnostic procedures including a full-field electroretinogram and the cortical-derived visual evoked potential (*65*) and were blinded to the infant’s inoculation group. The electroretinogram is used clinically to assess generalized retinal function under light-adapted (focused on cone photoreceptors) conditions (*65*). Visual evoked potentials reflect the function of the entire visual pathway from the retina via the optic nerve to the visual cortex of the brain (*65*). The light-adapted full-field flash electroretinogram, recorded on a rod saturating background, measures the electrical activity generated predominantly by cone photoreceptor and bipolar cells, which are found in high density in the primate macula and are primarily responsible for light-adapted, high acuity and color vision.

Binocular photopic (light-adapted) electroretinography flashes (2.5 cd·s·m^-2^ and 30-Hz flicker) followed by monocularly stimulated (right eye, then left eye) visual evoked cortical potentials were recorded and collected. Measurements were recorded using an LKC UTAS visual electrodiagnostic test system configured with the BigShot Ganzfeld stimulator for electroretinogram and visual evoked potentials response collection (LKC Technologies™, Gaithersburg, MD). When isoflurane was used during an earlier procedure in the same sedation event, a washout period was allowed before visual electrophysiology studies to minimize isoflurane suppression of cortical activity(*66*). For the electroretinogram, corneas were anesthetized with topical 0.5% proparacaine prior to application of ERG-Jet™ (Universo™, Switzerland) contact lens electrodes and a conductive wetting solution. Four-channel (channel 1: right eye, channel 2: left eye, channel 3: right occipital cortex, channel 4: left occipital cortex) recordings were carried out with the use of the UBA patient amplifier interfaced to the system computer. The UBA amplifier response was calibrated at the initiation of the study. Reference electrodes were subdermal stainless-steel needle electrodes inserted near the ipsilateral outer canthus of each eye. To measure visual evoked potentials, a light stimulus was applied to either the left or right eyes (the non-tested eye was covered with an opaque occluder), and electrical potentials were recorded near the occipital cortex from two active subdermal electrodes situated approximately 1 cm superior to the occipital ridge and 1 cm lateral to the midline; reference electrodes were situated along the midline at the vertex. ERG-Jet™ and subdermal needle electrode impedances were < 5 kΩ (Grass Electrode Impedance Meter Model F-EZM5, Astro-Med, Inc., Grass Instrument Division, West Warwick, RI) and equivalent. The band pass for electroretinogram signals was 0.3 Hz (high pass) and 500 Hz (low pass) and the band pass for visual evoked potentials was 1 Hz (high pass) and 100 Hz (low pass). Each of the two electroretinogram flash stimulations were repeated for replicability. Replicates (2–4) of 80 flash stimulations each were performed on each macaque for visual evoked potentials. Response waveforms were digitized at 0.5 msec intervals. All response waveforms were processed off-line. The root mean square of the visual evoked potential amplitude from 50-200 msec post flash was machine-scored using software written in MATLAB™ (Natick, MA).

### Statistical analyses

#### Study demographics

In order to better understand the infants included in this study, several statistical analyses were performed to obtain the demographic features of exposure groups. Descriptive statistics including means, confidence intervals, and missing data were determined for each exposure group for demographic features measured on a quantitative scale. Categorical features were summarized in terms of frequencies and percentages. Comparisons of demographic features between exposure groups were conducted using a two-sample t-test for quantitative variables and chi-square/Fisher’s exact test for categorical variables. A two-sided 0.05 significance level was used to determine statistical significance for all comparisons.

#### Developmental outcomes

Developmental outcomes were reported in the form of continuous quantitative data, discrete quantitative data, and ordinal and nominal qualitative data, as ratio/percentage, frequency, or categorical values respectively. Group-level summaries were generated separately for ZIKV-exposed and control animals. Categorical data for the sensory processing task were summarized in terms of frequencies and percentages (n (%)).

For continuous behavioral outcomes measured on a quantitative scale, including puzzle feeder metrics and sensory processing measures as well as all variables for the PVC and home cage observation coding schemes, we reported number of observations, medians, and interquartile ranges (IQRs) - . Comparisons between exposure groups between those outcomes were conducted using analysis of variance (ANOVA), followed by pairwise post-hoc comparisons. Chi-square or Fisher’s exact test was used to compare binary development outcomes between groups. Statistical inference was conducted using SAS software (SAS Institute, Cary NC), version 9.4.

#### Analyses for evaluating associations between infant weight, maternal, and placental virologic parameters

Linear regression models were used to assess associations between infant weight at 12 months and key maternal or placental virologic parameters, including maternal plasma viremia duration, area under the curve of plasma viral loads, proportion of maternal-fetal interface biopsies positive for viral RNA, and PRNT90 neutralizing antibody titers. Data were analyzed in R (version 4.x) using the lm() function. Each parameter was evaluated in a separate univariate model with infant weight as the outcome. Prior to modeling, variables were explicitly converted to numeric type, and records with missing values for either the predictor or outcome variable were excluded. Model fit was visualized using ggplot2, with scatter plots displaying individual data points and fitted regression lines. R² and p-values were reported for each model to quantify the strength and significance of the associations.

#### Mediation analyses

We conducted causal mediation analyses to evaluate whether the relationship between prenatal ZIKV exposure and 12-month behavioral outcomes was mediated by hearing loss or visual pathway function. Behavioral outcomes included ventral contact duration and feather approach behavior, which was assessed both as a 4-level ordinal variable and as a binary outcome (approached vs. did not approach). Predictor variables included plasma viremia duration, maternal PRNT90 neutralizing antibody titer at 1 month, ZIKV inoculation group, and binary exposure classification (ZIKV-exposed vs. control). For each predictor–mediator–outcome combination, we fitted two linear models: a mediator model (mediator ∼ predictor) and an outcome model (outcome ∼ predictor + mediator). Mediation analysis was performed using the mediate() function from the R mediation package (version 2024.12.0), with 1,000 simulations and quasi-Bayesian approximation for inference. All continuous outcomes and mediators were modeled using linear regression. Binary outcomes (feather approach) and binary mediators (hearing loss) were treated as numeric (0/1) and analyzed using linear models given the small sample size and model convergence considerations. Mediation significance was evaluated by examining the average causal mediation effect (ACME), average direct effect (ADE), total effect, and proportion mediated. All analyses were conducted using R version 2024.12.0.

### Study approval

The University of Wisconsin-Madison Institutional Animal Care and Use Committee approved the nonhuman primate research covered under protocol numbers G006290 and G006108.

## List of Supplementary Materials

Fig. S1 Fig. S2 Fig. S3 Fig. S4 Fig. S5 Fig. S6 Table S1 Table S2 Table S3 Table S4 Table S5 Table S6 Table S7 Table S8 Table S9 Table S10 Table S11 Table S12

## Acknowledgments

We thank the Wisconsin National Primate Research Center, specifically Behavioral Management Services, Scientific Protocol Implementation Services, Pathology Services, and Veterinary Services for their assistance with this project. We thank David O’Connor for insightful discussions on experimental design. This project was supported in part by NIH P01AI132132. We used ChatGPT 4o to assist with improving readability. We thank Saswati Bhattacharya, Taylor Treadway, and Nikunj Makwana for their contributions.

## Funding

National Institutes of Health grant R01 AI153130 (KKA, ELM)

National Institutes of Health grant P30EY016665 (Vision Research Core grant)

Eunice Kennedy Shriver National Institute of Child Health and Human Development P50HD105353 (Waisman Center)

## Author contributions

Conceptualization: KKA, ELM

Methodology: KKA, AH, AMW, CR, MN, HAS, JVH, TCF, ELM

Investigation: KKA, BB, ERR, JK, NPK, AMM, SW, VM, JRD, SK, FE, RVS, AS, AS, AK, CK, AH, AMW, CR, MN, PB, HAS, JVH, ELM

Visualization: ERR, AS, AS, JVH, ELM Funding acquisition: KKA, ELM Project administration: KKA, ELM

Supervision: KKA, AH, MN, HAS, SC, TCF Writing – original draft: KKA, BB, ERR, JK

Writing – review & editing: KKA, BB, ERR, JK, NPK, AMM, SW, VM, JRD, SK, FE, RVS, AS, AS, AK, CK, AH, AMW, CR, MN, PB, HAS, JVH, TCF, ELM

**Competing interests:** James Ver Hoeve is a consultant for a company called OSOD, A Merit Company, which provides consulting services to the pharmaceutical industry.

**Data and materials availability:** All raw data associated with this manuscript that is not presented within the main test or in supplementary files is available at: https://go.wisc.edu/i27a67.

## References and Notes

1. G. R. Deshpande, G. N. Sapkal, A. Salunke, R. Gunjikar, N. Tadkalkar, P. Shinde, N. Daga, M. Gopale, A. Ramdasi, S. Hundekar, K. Lole, R. R. Roy, J. A. Jenish, R. Srivastava, S. Parmar, P. Pawara, K. Jarande, S. Vidhate, K. Khutwad, An outbreak of Zika virus in western India in the metropolis of Pune in the monsoon of 2024. J. Infect. Public Health 18, 102720 (2025).

2. L. Pezzi, N. Ayhan, J. Brulé, G. A. Durand, G. Grard, X. Lamballerie, R. Klitting, Zika virus infection in a traveller returning to France from Seychelles, 2024. J. Travel Med., doi: 10.1093/jtm/taaf048 (2025).

3. S. B. Mulkey, M. Arroyave-Wessel, C. Peyton, D. I. Bulas, Y. Fourzali, J. Jiang, S. Russo, R. McCarter, M. E. Msall, A. J. du Plessis, R. L. DeBiasi, C. Cure, Neurodevelopmental Abnormalities in Children With In Utero Zika Virus Exposure Without Congenital Zika Syndrome. JAMA Pediatr. 174, 269–276 (2020).

4. K. Nielsen-Saines, P. Brasil, T. Kerin, Z. Vasconcelos, C. R. Gabaglia, L. Damasceno, M. Pone, L. M. Abreu de Carvalho, S. M. Pone, A. A. Zin, I. Tsui, T. R. S. Salles, D. C. da Cunha, R. P. Costa, J. Malacarne, A. B. Reis, R. H. Hasue, C. Y. P. Aizawa, F. F. Genovesi, C. Einspieler, P. B. Marschik, J. P. Pereira, S. L. Gaw, K. Adachi, J. D. Cherry, Z. Xu, G. Cheng, M. E. Moreira, Delayed childhood neurodevelopment and neurosensory alterations in the second year of life in a prospective cohort of ZIKV-exposed children. Nat. Med. 25, 1213–1217 (2019).

5. P. M. Peçanha, S. C. Gomes Junior, S. M. Pone, M. V. da S. Pone, Z. Vasconcelos, A. Zin, R. H. H. Vilibor, R. P. Costa, M. D. B. B. Meio, K. Nielsen-Saines, P. Brasil, E. Brickley, M. E. Lopes Moreira, Neurodevelopment of children exposed intra-uterus by Zika virus: A case series. PLoS One 15, e0229434 (2020).

6. R. A. de O. Vianna, K. L. Lovero, S. A. de Oliveira, A. R. Fernandes, T. C. S. D. Santos, L. C. S. De S. Lima, F. R. Carvalho, M. D. S. Quintans, A. C. Bueno, A. F. M. Torbey, A. L. A. A. G. de Souza, A. de O. P. de Farias, L. A. B. Camacho, L. W. Riley, C. A. A. Cardoso, Children Born to Mothers with Rash During Zika Virus Epidemic in Brazil: First 18 Months of Life. J. Trop. Pediatr. 65, 592– 602 (2019).

7. F. A. Venancio, M. E. Quilião, S. C. de O. Gabeira, A. T. de Carvalho, S. H. D. S. Leite, S. M. B. de Lima, N. D. S. Alves, L. da C. Moura, W. D. Schwarcz, A. de S. Azevedo, L. H. F. Demarchi, M. C. S. U. Zardin, G. G. de C. Lichs, D. L. Taira, W. de S. Fernandes, N. O. Alves, A. E. C. Arrua, A. I. do Nascimento, L. K. Mareto, M. V. de Azevedo, C. G. Maciel, M. J. de Medeiros, M. M. de S. Rodrigues, Z. Vasconcelos, K. Nielsen-Saines, R. V. da Cunha, C. D. B. Santos-Pinto, E. F. de Oliveira, Early and long-term adverse outcomes of in utero Zika exposure. Pediatrics 155 (2025).

8. S. B. Mulkey, R. Andringa-Seed, E. Corn, M. E. Williams, M. Arroyave-Wessel, R. H. Podolsky, C. Peyton, M. E. Msall, C. Cure, M. M. Berl, School-age child neurodevelopment following antenatal Zika virus exposure. Pediatr. Res., doi: 10.1038/s41390-025-03981-7 (2025).

9. C. A. Moore, J. E. Staples, W. B. Dobyns, A. Pessoa, C. V. Ventura, E. B. da Fonseca, E. M. Ribeiro, L. O. Ventura, N. N. Neto, J. F. Arena, S. A. Rasmussen, Characterizing the Pattern of Anomalies in Congenital Zika Syndrome for Pediatric Clinicians. JAMA Pediatr. 171, 288–295 (2017).

10. N. M. Roth, M. R. Reynolds, E. L. Lewis, K. R. Woodworth, S. Godfred-Cato, A. Delaney, A. Akosa, M. Valencia-Prado, M. Lash, A. Elmore, P. Langlois, S. Khuwaja, A. Tufa, E. M. Ellis, E. Nestoridi, C. Lyu, N. D. Longcore, M. Piccardi, L. Lind, S. Starr, L. Johnson, S. E. Browne, M. Gosciminski, P. E. Velasco, F. Johnson-Clarke, A. Locklear, M. Chan, J. Fornoff, K.-A. E. Toews, J. Tonzel, N. S. Marzec, S. Hale, A. E. Nance, T. Willabus, D. Contreras, S. N. Adibhatla, L. Iguchi, E. Potts, E. Schiffman, K. Lolley, B. Stricklin, E. Ludwig, H. Garstang, M. Marx, E. Ferrell, C. Moreno-Gorrin, K. Signs, P. Romitti, V. Leedom, B. Martin, L. Castrodale, A. Cook, C. Fredette, L. Denson, L. Cronquist, J. F. Nahabedian 3rd, N. Shinde, K. Polen, S. M. Gilboa, S. W. Martin, J. D. Cragan, D. Meaney-Delman, M. A. Honein, V. T. Tong, C. A. Moore, Zika-Associated Birth Defects Reported in Pregnancies with Laboratory Evidence of Confirmed or Possible Zika Virus Infection - U.S. Zika Pregnancy and Infant Registry, December 1, 2015-March 31, 2018. MMWR Morb. Mortal. Wkly. Rep. 71, 73–79 (2022).

11. A. Mahmoud, L. Pomar, V. Lambert, O. Picone, N. Hcini, Prenatal and Postnatal Ocular Abnormalities Following Congenital Zika Virus Infections: A Systematic Review. Ocul. Immunol. Inflamm., 1–11 (2024).

12. L. C. de Almeida, L. F. Muniz, R. J. Maciel, D. S. Ramos, K. M. G. de Albuquerque, Â. M. C. Leão, M. V. de Mendonça, M. de C. Leal, Hearing and communicative skills in the first years of life in children with congenital Zika syndrome. Braz. J. Otorhinolaryngol. 88, 112–117 (2022).

13. M. H. de M. Barbosa, M. C. de Magalhães-Barbosa, J. R. Robaina, A. Prata-Barbosa, M. A. de M. T. de Lima, A. J. L. A. da Cunha, Auditory findings associated with Zika virus infection: an integrative review. Braz. J. Otorhinolaryngol. 85, 642–663 (2019).

14. C. Veldhorst, M. Vervloed, S. Kef, B. Steenbergen, A scoping review of longitudinal studies of children with vision impairment. Br. J. Vis. Impair. 41, 587–609 (2023).

15. J. E. C. Lieu, M. Kenna, S. Anne, L. Davidson, Hearing loss in children: A review: A review. JAMA 324, 2195–2205 (2020).

16. E. L. Mohr, Modeling Zika Virus-Associated Birth Defects in Nonhuman Primates. J Pediatric Infect Dis Soc 7, S60–S66 (2018).

17. D. M. Dudley, M. T. Aliota, E. L. Mohr, C. M. Newman, T. G. Golos, T. C. Friedrich, D. H. O’Connor, Using Macaques to Address Critical Questions in Zika Virus Research. Annu Rev Virol 6, 481–500 (2019).

18. H. Narasimhan, A. Chudnovets, I. Burd, A. Pekosz, S. L. Klein, Animal models of congenital zika syndrome provide mechanistic insight into viral pathogenesis during pregnancy. PLoS Negl. Trop. Dis. 14, e0008707 (2020).

19. T. E. Morrison, M. S. Diamond, Animal Models of Zika Virus Infection, Pathogenesis, and Immunity. J. Virol. 91 (2017).

20. J. Gutkes, N. P. Krabbe, K. Ausderau, E. L. Mohr, Macaque models of prenatal and postnatal Zika virus exposure and developmental outcomes. J. Pediatric Infect. Dis. Soc., doi: 10.1093/jpids/piaf024 (2025).

21. G. P. Sackett, “CHAPTER 1 - Developmental Disabilities and Primate Models Defined” in Primate Models of Children’s Health and Developmental Disabilitie*s*, T. M. Burbacher, G. P. Sackett, K. S. Grant, Eds. (Academic Press, New York, 2008), pp. 1–10.

22. M. R. Koenig, E. Razo, A. Mitzey, C. M. Newman, D. M. Dudley, M. E. Breitbach, M. R. Semler, L. M. Stewart, A. M. Weiler, S. Rybarczyk, K. M. Bach, M. S. Mohns, H. A. Simmons, A. Mejia, M. Fritsch, M. Dennis, L. B. C. Teixeira, M. L. Schotzko, T. M. Nork, C. A. Rasmussen, A. Katz, V. Nair, J. Hou, A. Hartman, J. Ver Hoeve, C. Kim, M. L. Schneider, K. Ausderau, S. Kohn, A. S. Jaeger, M. T. Aliota, J. M. Hayes, N. Schultz-Darken, J. Eickhoff, K. M. Antony, K. Noguchi, X. Zeng, S. Permar, V. Prabhakaran, S. Capuano 3rd, T. C. Friedrich, T. G. Golos, D. H. O’Connor, E. L. Mohr, Quantitative definition of neurobehavior, vision, hearing and brain volumes in macaques congenitally exposed to Zika virus. PLoS One 15, e0235877 (2020).

23. K. Ausderau, S. Kabakov, E. Razo, A. M. Mitzey, K. M. Bach, C. M. Crooks, N. Dulaney, L. Keding, C. Salas-Quinchucua, L. G. Medina-Magües, A. M. Weiler, M. Bliss, J. Eickhoff, H. A. Simmons, A. Mejia, K. M. Antony, T. Morgan, S. Capuano 3rd, M. L. Schneider, M. T. Aliota, T. C. Friedrich, D. H. O’Connor, T. G. Golos, E. L. Mohr, Neonatal Development in Prenatally Zika Virus-Exposed Infant Macaques with Dengue Immunity. Viruses 13 (2021).

24. L. Pomar, V. Lambert, S. Matheus, C. Pomar, N. Hcini, G. Carles, D. Rousset, M. Vouga, A. Panchaud, D. Baud, Prolonged Maternal Zika Viremia as a Marker of Adverse Perinatal Outcomes. Emerg. Infect. Dis. 27, 490–498 (2021).

25. R. W. Driggers, C.-Y. Ho, E. M. Korhonen, S. Kuivanen, A. J. Jääskeläinen, T. Smura, A. Rosenberg, D. A. Hill, R. L. DeBiasi, G. Vezina, J. Timofeev, F. J. Rodriguez, L. Levanov, J. Razak, P. Iyengar, A. Hennenfent, R. Kennedy, R. Lanciotti, A. du Plessis, O. Vapalahti, Zika Virus Infection with Prolonged Maternal Viremia and Fetal Brain Abnormalities. N. Engl. J. Med. 374, 2142–2151 (2016).

26. A. Suy, E. Sulleiro, C. Rodó, É. Vázquez, C. Bocanegra, I. Molina, J. Esperalba, M. P. Sánchez-Seco, H. Boix, T. Pumarola, E. Carreras, Prolonged Zika Virus Viremia during Pregnancy. N. Engl. J. Med. 375, 2611–2613 (2016).

27. K. L. Schwartz, T. Chan, N. Rai, K. E. Murphy, W. Whittle, M. A. Drebot, J. Gubbay, A. K. Boggild, Zika virus infection in a pregnant Canadian traveler with congenital fetal malformations noted by ultrasonography at 14-weeks gestation. Trop Dis Travel Med Vaccines 4, 2 (2018).

28. K. Nielsen-Saines, T. Kalbasi-Romero, A. C. M. Duarte, S. Almeida da Silva, K. Adachi, L. Damasceno, T. Kerin, T. Fuller, J. G. Deville, M. E. Moreira, Z. Vasconcelos, A. Zin, M. Shin-Sim, S. M. Barbosa de Lima, P. Brasil, Development of maternal antibodies post ZIKV in pregnancy is associated with lower risk of microcephaly and structural brain abnormalities in exposed infants. J. Infect. Dis., doi: 10.1093/infdis/jiaf146 (2025).

29. A. Gordon, L. Gresh, S. Ojeda, L. C. Katzelnick, N. Sanchez, J. C. Mercado, G. Chowell, B. Lopez, D. Elizondo, J. Coloma, R. Burger-Calderon, G. Kuan, A. Balmaseda, E. Harris, Prior dengue virus infection and risk of Zika: A pediatric cohort in Nicaragua. PLoS Med. 16, e1002726 (2019).

30. S. V. Bardina, P. Bunduc, S. Tripathi, J. Duehr, J. J. Frere, J. A. Brown, R. Nachbagauer, G. A. Foster, D. Krysztof, D. Tortorella, S. L. Stramer, A. García-Sastre, F. Krammer, J. K. Lim, Enhancement of Zika virus pathogenesis by preexisting antiflavivirus immunity. Science 356, 175– 180 (2017).

31. T. Langerak, M. Broekhuizen, P.-P. A. Unger, L. Tan, M. Koopmans, E. van Gorp, A. H. J. Danser, B. Rockx, Transplacental Zika virus transmission in ex vivo perfused human placentas. PLoS Negl. Trop. Dis. 16, e0010359 (2022).

32. M. K. McCracken, G. D. Gromowski, H. L. Friberg, X. Lin, P. Abbink, R. De La Barrera, K. H. Eckles, L. S. Garver, M. Boyd, D. Jetton, D. H. Barouch, M. C. Wise, B. S. Lewis, J. R. Currier, K. Modjarrad, M. Milazzo, M. Liu, A. B. Mullins, J. R. Putnak, N. L. Michael, R. G. Jarman, S. J. Thomas, Impact of prior flavivirus immunity on Zika virus infection in rhesus macaques. PLoS Pathog. 13, e1006487 (2017).

33. P. Pantoja, E. X. Pérez-Guzmán, I. V. Rodríguez, L. J. White, O. González, C. Serrano, L. Giavedoni, V. Hodara, L. Cruz, T. Arana, M. I. Martínez, M. A. Hassert, J. D. Brien, A. K. Pinto, A. de Silva, C. A. Sariol, Zika virus pathogenesis in rhesus macaques is unaffected by pre-existing immunity to dengue virus. Nat. Commun. 8, 15674 (2017).

34. N. M. S. Sansone, M. N. Boschiero, F. A. L. Marson, Dengue outbreaks in Brazil and Latin America: the new and continuing challenges. Int. J. Infect. Dis. 147, 107192 (2024).

35. U.-A. Halai, K. Nielsen-Saines, M. L. Moreira, P. C. de Sequeira, J. P. P. Junior, A. de Araujo Zin, J. Cherry, C. R. Gabaglia, S. L. Gaw, K. Adachi, I. Tsui, J. H. Pilotto, R. R. Nogueira, A. M. B. de Filippis, P. Brasil, Maternal Zika Virus Disease Severity, Virus Load, Prior Dengue Antibodies, and Their Relationship to Birth Outcomes. [Preprint] (2017). 10.1093/cid/cix472.

36. P. Brasil, J. P. Pereira Jr, M. E. Moreira, R. M. Ribeiro Nogueira, L. Damasceno, M. Wakimoto, R. S. Rabello, S. G. Valderramos, U.-A. Halai, T. S. Salles, A. A. Zin, D. Horovitz, P. Daltro, M. Boechat, C. Raja Gabaglia, P. Carvalho de Sequeira, J. H. Pilotto, R. Medialdea-Carrera, D. Cotrim da Cunha, L. M. Abreu de Carvalho, M. Pone, A. Machado Siqueira, G. A. Calvet, A. E. Rodrigues Baião, E. S. Neves, P. R. Nassar de Carvalho, R. H. Hasue, P. B. Marschik, C. Einspieler, C. Janzen, J. D. Cherry, A. M. Bispo de Filippis, K. Nielsen-Saines, Zika Virus Infection in Pregnant Women in Rio de Janeiro. N. Engl. J. Med. 375, 2321–2334 (2016).

37. A. Moreira-Soto, R. Cabral, C. Pedroso, M. Eschbach-Bludau, A. Rockstroh, L. A. Vargas, I. Postigo-Hidalgo, E. Luz, G. S. Sampaio, C. Drosten, E. M. Netto, T. Jaenisch, S. Ulbert, M. Sarno, C. Brites, J. F. Drexler, Exhaustive TORCH Pathogen Diagnostics Corroborate Zika Virus Etiology of Congenital Malformations in Northeastern Brazil. mSphere 3 (2018).

38. C. Pedroso, C. Fischer, M. Feldmann, M. Sarno, E. Luz, A. Moreira-Soto, R. Cabral, E. M. Netto, C. Brites, B. M. Kümmerer, J. F. Drexler, Cross-protection of dengue virus infection against congenital Zika syndrome, northeastern Brazil. Emerg. Infect. Dis. 25, 1485–1493 (2019).

39. B. de Paula Freitas, J. R. de Oliveira Dias, J. Prazeres, G. A. Sacramento, A. I. Ko, M. Maia, R. Belfort Jr, Ocular Findings in Infants With Microcephaly Associated With Presumed Zika Virus Congenital Infection in Salvador, Brazil. JAMA Ophthalmol. 134, 529–535 (2016).

40. R. Pimentel, S. Khosla, J. Rondon, F. Peña, G. Sullivan, M. Perez, S. D. Mehta, M. O. Brito, Birth Defects and Long-Term Neurodevelopmental Abnormalities in Infants Born During the Zika Virus Epidemic in the Dominican Republic. Ann Glob Health 87, 4 (2021).

41. M. A. Honein, A. L. Dawson, E. E. Petersen, A. M. Jones, E. H. Lee, M. M. Yazdy, N. Ahmad, J. Macdonald, N. Evert, A. Bingham, S. R. Ellington, C. K. Shapiro-Mendoza, T. Oduyebo, A. D. Fine, C. M. Brown, J. N. Sommer, J. Gupta, P. Cavicchia, S. Slavinski, J. L. White, S. M. Owen, L. R. Petersen, C. Boyle, D. Meaney-Delman, D. J. Jamieson, US Zika Pregnancy Registry Collaboration, Birth Defects Among Fetuses and Infants of US Women With Evidence of Possible Zika Virus Infection During Pregnancy. JAMA 317, 59–68 (2017).

42. A. S. Jaeger, R. A. Murrieta, L. R. Goren, C. M. Crooks, R. V. Moriarty, A. M. Weiler, S. Rybarczyk, M. R. Semler, C. Huffman, A. Mejia, H. A. Simmons, M. Fritsch, J. E. Osorio, J. C. Eickhoff, S. L. O’Connor, G. D. Ebel, T. C. Friedrich, M. T. Aliota, Zika viruses of African and Asian lineages cause fetal harm in a mouse model of vertical transmission. PLoS Negl. Trop. Dis. 13, e0007343 (2019).

43. C. M. Crooks, A. M. Weiler, S. L. Rybarczyk, M. Bliss, A. S. Jaeger, M. E. Murphy, H. A. Simmons, A. Mejia, M. K. Fritsch, J. M. Hayes, J. C. Eickhoff, A. M. Mitzey, E. Razo, K. M. Braun, E. A. Brown, K. Yamamoto, P. M. Shepherd, A. Possell, K. Weaver, K. M. Antony, T. K. Morgan, X. Zeng, D. M. Dudley, E. Peterson, N. Schultz-Darken, D. H. O’Connor, E. L. Mohr, T. G. Golos, M. T. Aliota, T. C. Friedrich, African-Lineage Zika Virus Replication Dynamics and Maternal-Fetal Interface Infection in Pregnant Rhesus Macaques. J. Virol. 95, e0222020 (2021).

44. L. E. Raasch, K. Yamamoto, C. M. Newman, J. R. Rosinski, P. M. Shepherd, E. Razo, C. M. Crooks, M. I. Bliss, M. E. Breitbach, E. L. Sneed, A. M. Weiler, X. Zeng, K. K. Noguchi, T. K. Morgan, N. A. Fuhler, E. K. Bohm, A. J. Alberts, S. J. Havlicek, S. Kabakov, A. M. Mitzey, K. M. Antony, K. K. Ausderau, A. Mejia, P. Basu, H. A. Simmons, J. C. Eickhoff, M. T. Aliota, E. L. Mohr, T. C. Friedrich, T. G. Golos, D. H. O’Connor, D. M. Dudley, Fetal loss in pregnant rhesus macaques infected with high-dose African-lineage Zika virus. PLoS Negl. Trop. Dis. 16, e0010623 (2022).

45. E. L. Mohr, L. N. Block, C. M. Newman, L. M. Stewart, M. Koenig, M. Semler, M. E. Breitbach, L. B. C. Teixeira, X. Zeng, A. M. Weiler, G. L. Barry, T. H. Thoong, G. J. Wiepz, D. M. Dudley, H. A. Simmons, A. Mejia, T. K. Morgan, M. S. Salamat, S. Kohn, K. M. Antony, M. T. Aliota, M. S. Mohns, J. M. Hayes, N. Schultz-Darken, M. L. Schotzko, E. Peterson, S. Capuano 3rd, J. E. Osorio, S. L. O’Connor, T. C. Friedrich, D. H. O’Connor, T. G. Golos, Ocular and uteroplacental pathology in a macaque pregnancy with congenital Zika virus infection. PLoS One 13, e0190617 (2018).

46. M. D. Bauman, A.-M. Iosif, S. E. P. Smith, C. Bregere, D. G. Amaral, P. H. Patterson, Activation of the maternal immune system during pregnancy alters behavioral development of rhesus monkey offspring. Biol. Psychiatry 75, 332–341 (2014).

47. J. Raper, Z. Kovacs-Balint, M. Mavigner, S. Gumber, M. W. Burke, J. Habib, C. Mattingly, D. Fair, E. Earl, E. Feczko, M. Styner, S. M. Jean, J. K. Cohen, M. S. Suthar, M. M. Sanchez, M. C. Alvarado, A. Chahroudi, Long-term alterations in brain and behavior after postnatal Zika virus infection in infant macaques. Nat. Commun. 11, 2534 (2020).

48. M. L. Schneider, C. F. Moore, L. L. Gajewski, J. A. Larson, A. D. Roberts, A. K. Converse, O. T. DeJesus, Sensory processing disorder in a primate model: evidence from a longitudinal study of prenatal alcohol and prenatal stress effects. Child Dev. 79, 100–113 (2008).

49. C. F. Moore, L. L. Gajewski, N. K. Laughlin, M. L. Luck, J. A. Larson, M. L. Schneider, Developmental lead exposure induces tactile defensiveness in rhesus monkeys (Macaca mulatta). Environ. Health Perspect. 116, 1322–1326 (2008).

50. P. R. Cogo, G. Moadab, E. Bliss-Moreau, F. Pittet, Prenatal Zika virus exposure alters the interaction between affective processing and decision-making in juvenile rhesus macaques (Macaca mulatta). Dev. Psychobiol. 66, e70002 (2024).

51. L. K. White, J. M. McDermott, K. A. Degnan, H. A. Henderson, N. A. Fox, Behavioral inhibition and anxiety: the moderating roles of inhibitory control and attention shifting. J. Abnorm. Child Psychol. 39, 735–747 (2011).

52. L. F. B. Almeida, M. Kattah, L. O. Ventura, A. L. Gois, C. Rocha, C. G. Andrade, C. Mendonza- Santiesteban, C. V. Ventura, Pattern-Reversal Visual Evoked Potential in Children With Congenital Zika Syndrome. J. Pediatr. Ophthalmol. Strabismus 58, 78–83 (2021).

53. M. García-Boyano, R. García-Segovia, A. Fernández-Menéndez, Y. Pérez, J. Bustamante-Amador, M. Layana-Coronel, J. M. Caballero-Caballero, C. Rodríguez-Izquierdo, N. Chávez-Solórzano, D. Solís-Montiel, G. Miño-León, Long-term outcomes of infants with congenital Zika virus infection in Ecuador: A retrospective longitudinal study. J. Trop. Pediatr. 67 (2021).

54. S. Gordon, A. Kerr, C. Wiggs, M. F. Chiang, What is cerebral/cortical visual impairment and why do we need a new definition? Ophthalmology 131, 1357–1358 (2024).

55. C. M. Crooks, A. M. Weiler, S. L. Rybarczyk, M. I. Bliss, A. S. Jaeger, M. E. Murphy, H. A. Simmons, A. Mejia, M. K. Fritsch, J. M. Hayes, J. C. Eickhoff, A. M. Mitzey, E. Razo, K. M. Braun, E. A. Brown, K. Yamamoto, P. M. Shepherd, A. Possell, K. Weaver, K. M. Antony, T. K. Morgan, C. M. Newman, D. M. Dudley, N. Schultz-Darken, E. Peterson, L. C. Katzelnick, A. Balmaseda, E. Harris, D. H. O’Connor, E. L. Mohr, T. G. Golos, T. C. Friedrich, M. T. Aliota, Previous exposure to dengue virus is associated with increased Zika virus burden at the maternal-fetal interface in rhesus macaques. [Preprint] (2021). 10.1371/journal.pntd.0009641.

56. C. L. Coe, G. R. Lubach, Maternal determinants of gestation length in the rhesus monkey. Trends Dev. Biol. 14, 63–72 (2021).

57. The Weatherall report on the use of non-human primates in research. https://royalsociety.org/topics-policy/publications/2006/weatherall-report.

58. National Research Council, Division on Earth and Life Studies, Institute for Laboratory Animal Research, Committee on Guidelines for the Use of Animals in Neuroscience and Behavioral Research, *Guidelines for the Care and Use of Mammals in Neuroscience and Behavioral Research* (National Academies Press, 2003).

59. A. F. Tarantal, Ultrasound Imaging in Rhesus (Macaca mulatta) and Long-tailed (Macaca fascicularis) Macaques: Reproductive and Research Applications. [Preprint] (2005). 10.1016/b978-012080261-6/50020-9.

60. D. M. Dudley, M. T. Alioto, E. L. Mohr, A. M. Weiler, G. Lehrer-Brey, K. L. Weisgrau, M. S. Mohns, M. E. Breitbach, M. N. Rasheed, C. M. Newman, D. D. Gellerup, L. H. Moncla, J. Post, N. Schultz-Darken, M. L. Schotzko, J. M. Hayes, J. A. Eudailey, M. A. Moody, S. R. Permar, S. L. O’Connor, E. G. Rakasz, H. A. Simmons, S. Capuano, T. G. Golos, J. E. Osorio, T. C. Friedrich, D. H. O’Connor, A rhesus macaque model of Asian-lineage Zika virus infection. Nat. Commun. 7, 12204 (2016).

61. S. G. Hansen, M. Piatak Jr, A. B. Ventura, C. M. Hughes, R. M. Gilbride, J. C. Ford, K. Oswald, R. Shoemaker, Y. Li, M. S. Lewis, A. N. Gilliam, G. Xu, N. Whizin, B. J. Burwitz, S. L. Planer, J. M. Turner, A. W. Legasse, M. K. Axthelm, J. A. Nelson, K. Früh, J. B. Sacha, J. D. Estes, B. F. Keele, P. T. Edlefsen, J. D. Lifson, L. J. Picker, Immune clearance of highly pathogenic SIV infection. Nature 502, 100–104 (2013).

62. N. P. Krabbe, E. Razo, H. J. Abraham, R. V. Spanton, Y. Shi, S. Bhattacharya, E. K. Bohm, J. C. Pritchard, A. M. Weiler, A. M. Mitzey, J. C. Eickhoff, E. Sullivan, J. C. Tan, M. T. Aliota, T. C. Friedrich, D. H. O’Connor, T. G. Golos, E. L. Mohr, Control of maternal Zika virus infection during pregnancy is associated with lower antibody titers in a macaque model. Front. Immunol. 14, 1267638 (2023).

63. J. W. Hall, New Handbook of Auditory Evoked Responses (Pearson, 2007).

64. Y. Huang, R. P. Danis, J. W. Pak, S. Luo, J. White, X. Zhang, A. Narkar, A. Domalpally, Development of a semi-automatic segmentation method for retinal OCT images tested in patients with diabetic macular edema. PLoS One 8, e82922 (2013).

65. A. G. Robson, J. Nilsson, S. Li, S. Jalali, A. B. Fulton, A. P. Tormene, G. E. Holder, S. E. Brodie, ISCEV guide to visual electrodiagnostic procedures. Doc. Ophthalmol. 136, 1–26 (2018).

66. G. Sitdikova, A. Zakharov, S. Janackova, E. Gerasimova, J. Lebedeva, A. R. Inacio, D. Zaynutdinova, M. Minlebaev, G. L. Holmes, R. Khazipov, Isoflurane suppresses early cortical activity. Ann Clin Transl Neurol 1, 15–26 (2014).

