## Supplementary materials for "Prenatal Zika Virus Exposure Disrupts Social-Emotional Development and Cortical Visual Function in Infant Macaques"

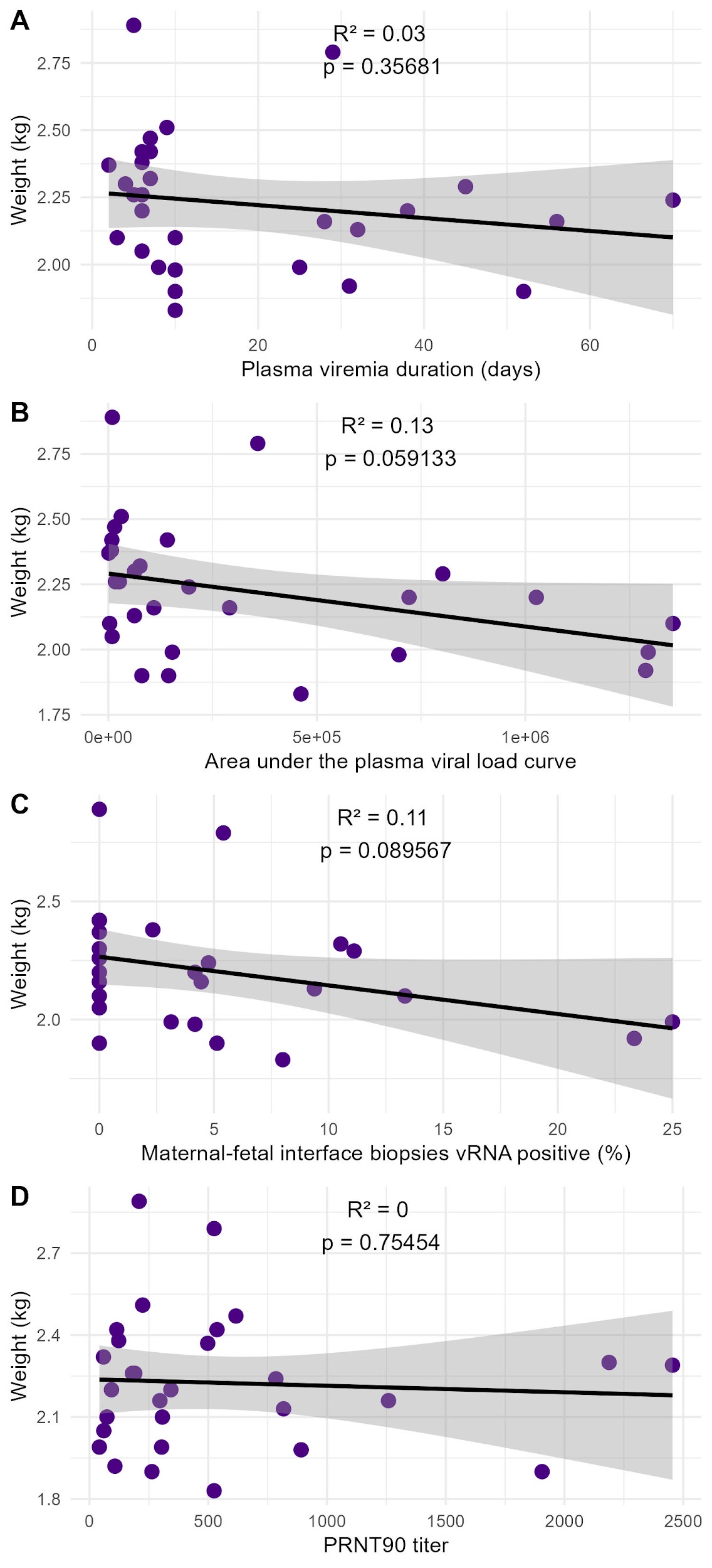

**Supplemental Figure 1.** ZIKV-exposed infant weight at 12 months of age and maternal viral and immunologic parameters. ZIKV-exposed infant weights at 12 months of age were compared to individual maternal variables plasma viremia duration (A), area under the curve plasma viral load (B), maternal-fetal interface biopsies vRNA positive (%) (C), and PRNT90 titer (D) using linear regression models.

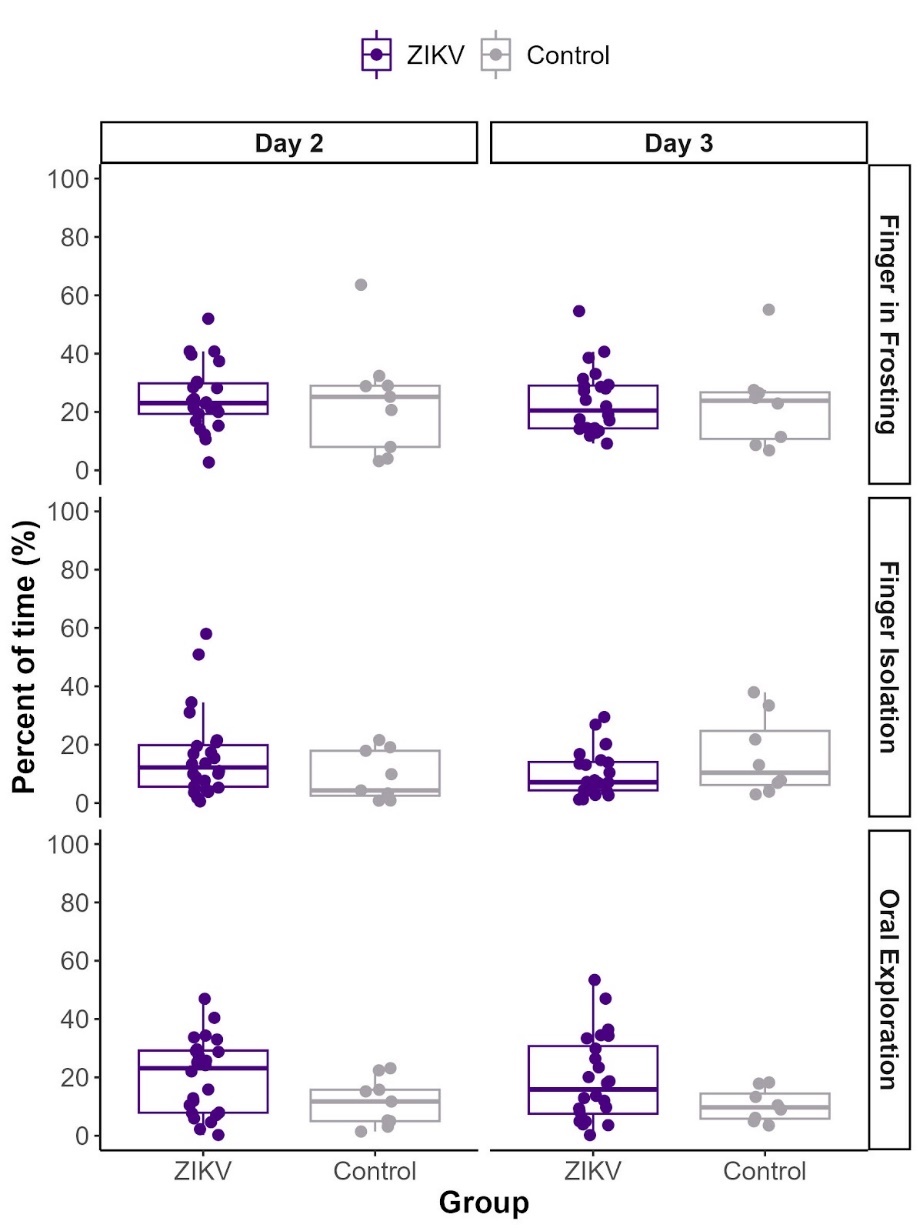

**Supplemental Figure 2**. PVC Pipe task evaluation of fine motor skills and sensory responsiveness. There were no significant differences between ZIKV-exposed and control infants in interactions with the PVC pipe, in the specific measurements of Finger in Frosting, Finger Isolation, and Oral Exploration.

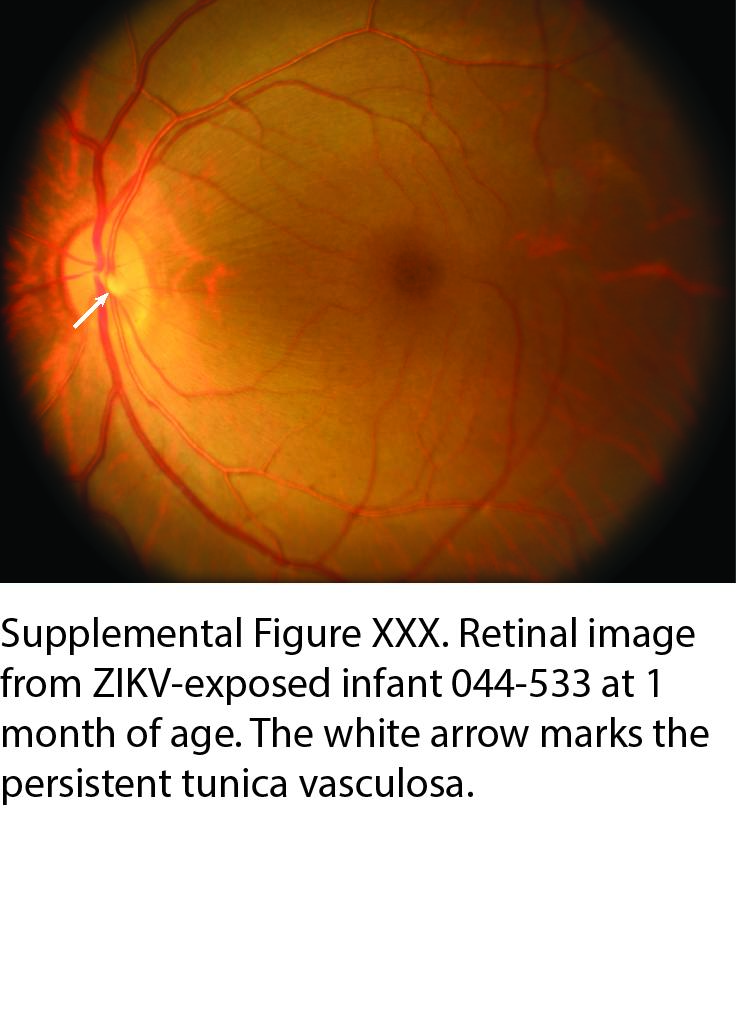

**Supplemental Figure 3.** Retinal image of ZIKV-exposed infant 044-533 showing persistent tunica vasculosa (white arrow).

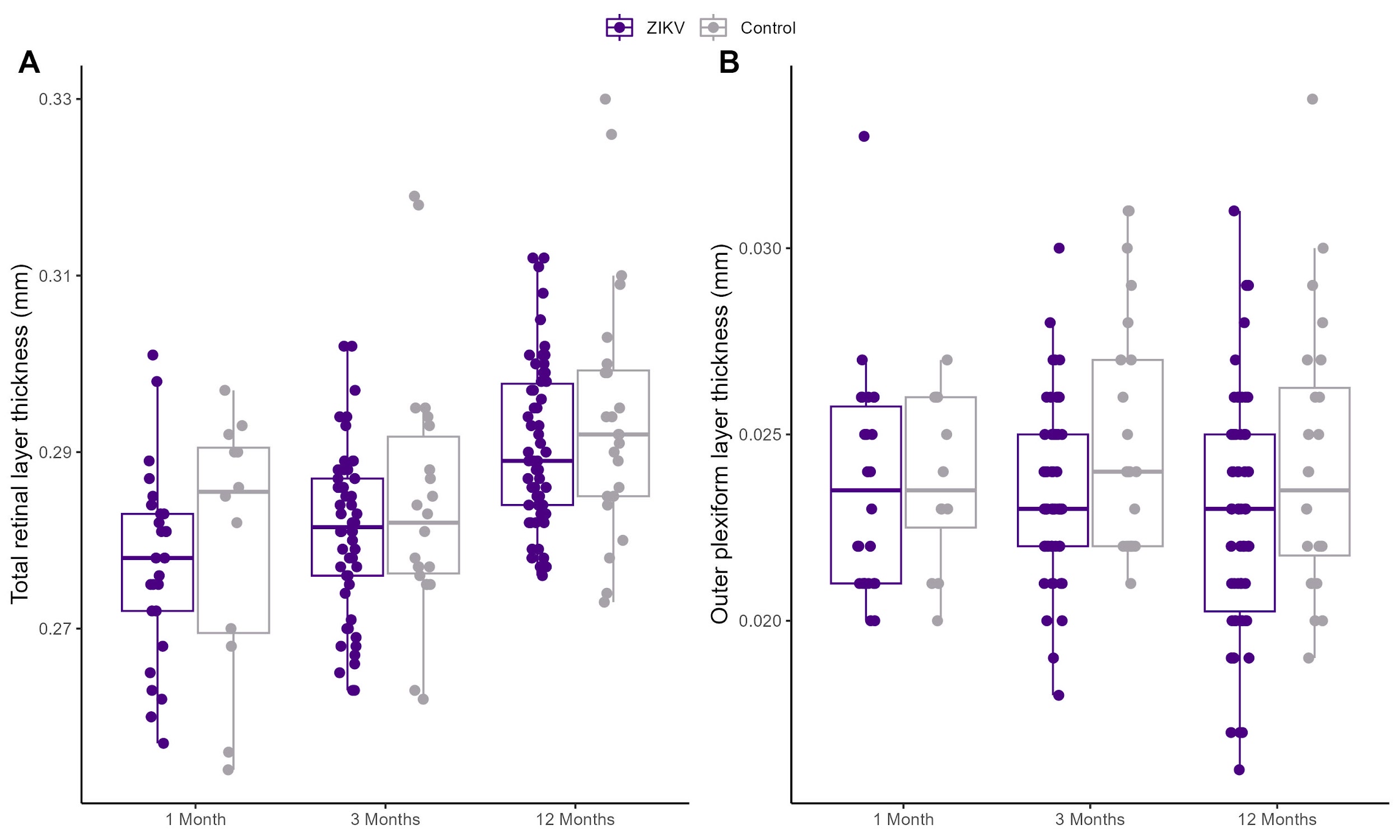

**Supplemental Figure 4.** Retinal layer thicknesses for ZIKV-exposed and control infants.

Retinal layer (A) and outer plexiform layer (B) thickness were obtained using spectral-domain optical coherence tomography and segmentation. The outer plexiform layer for ZIKV-exposed infants at 3 months and 12 months was slightly but not significantly lower than control infants (p = 0.0787 at 3 months; p = 0.0767 at 12 months). The right and left eye measurements are represented by individual points overlying the boxplots. Box plots show the interquartile range within the box, the median as a dark horizontal line, and the minimum and maximum values excluding the outliers are shown as whiskers.

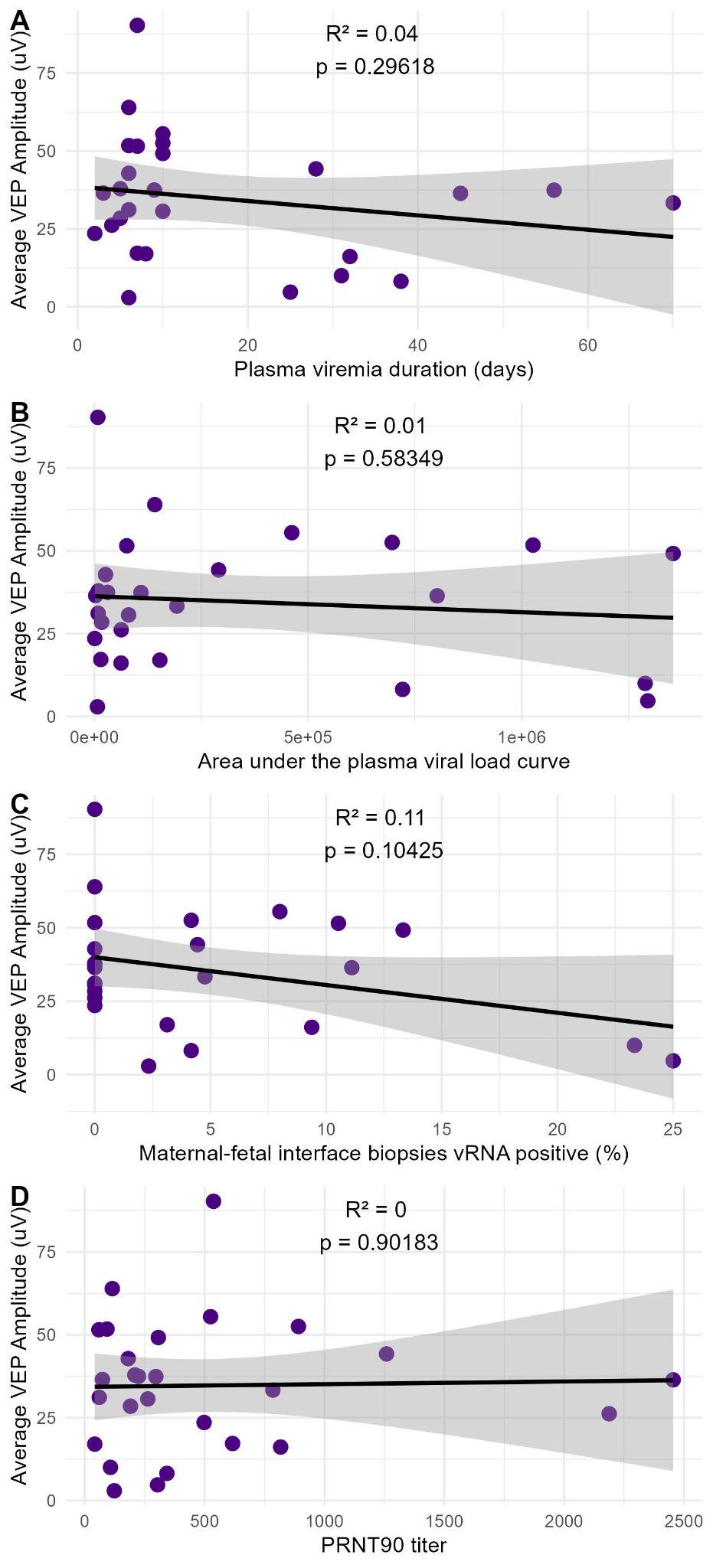

**Supplemental Figure 5.** Relationships between visual electrophysiology and maternal viral and immunologic parameters. There were no significant associations between visual electrophysiology amplitude at 3 months of age for ZIKV-exposed infants (with left and right sides averaged) and plasma viremia duration (A), area under the curve plasma viral load (B), maternal-fetal interface biopsies vRNA positive (%) (C), and PRNT90 titer, as defined with a linear regression.

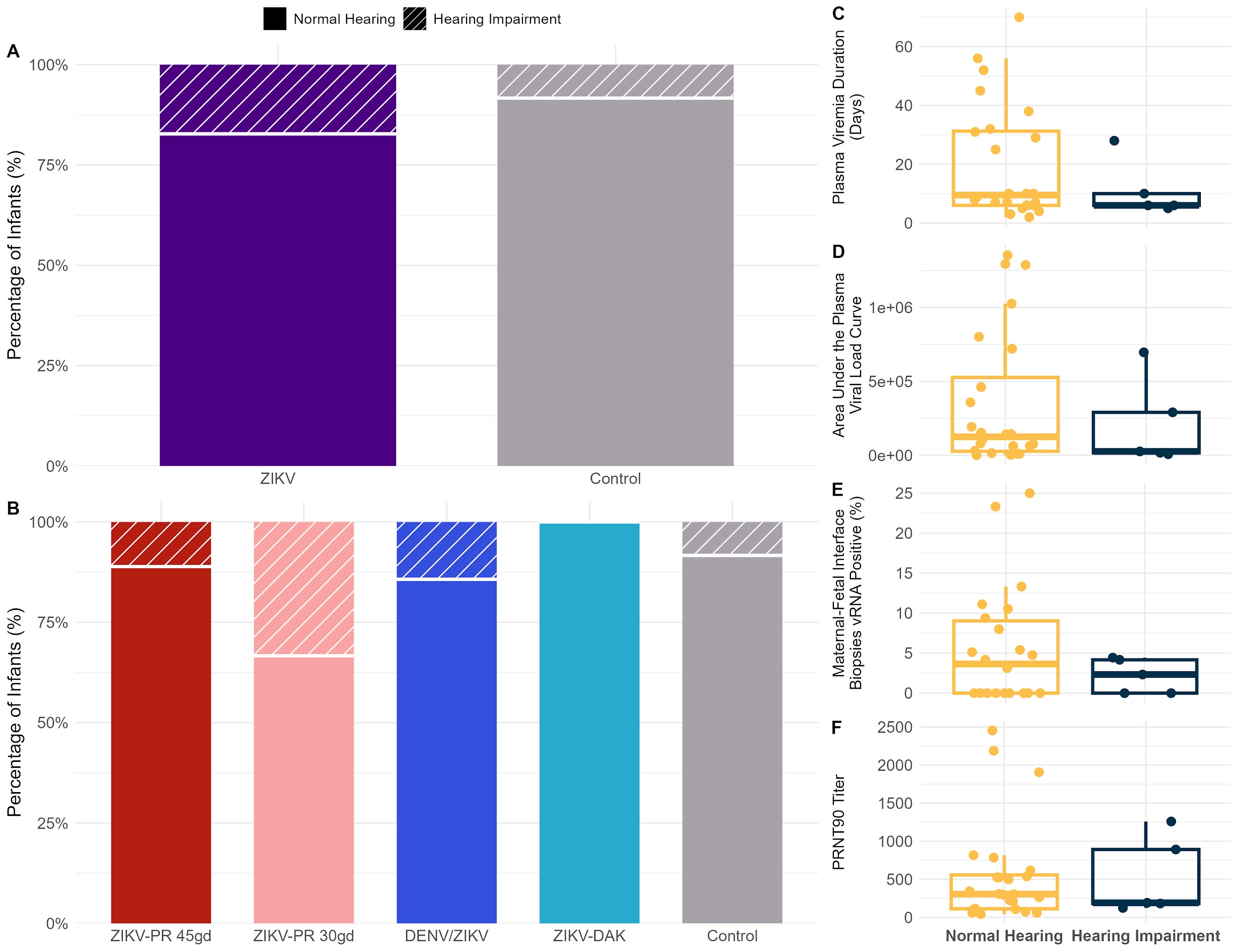

**Supplemental Figure 6.** Hearing loss in ZIKV-exposed and control infants. Auditory brain response test was performed with three stimuli (1000 Hz, 500 Hz, clicks) and a hearing loss was defined as the absence of Wave IV, i.e. no response, to any of the stimuli at the lowest volume tested. (A) Hearing loss occurred at a higher, but not significantly higher rate, in ZIKV-exposed infants compared to controls. (B) All of the ZIKV subgroups except for ZIKV-DAK had at least one infant with hearing loss. (C) There were no differences in maternal viral and immunologic parameters (plasma viremia duration (C), area under the curve plasma viral load (D), maternal-fetal interface biopsies that were vRNA positive (E) and PRNT90 titer (F) between the ZIKV-exposed infants with normal hearing and those with hearing loss.

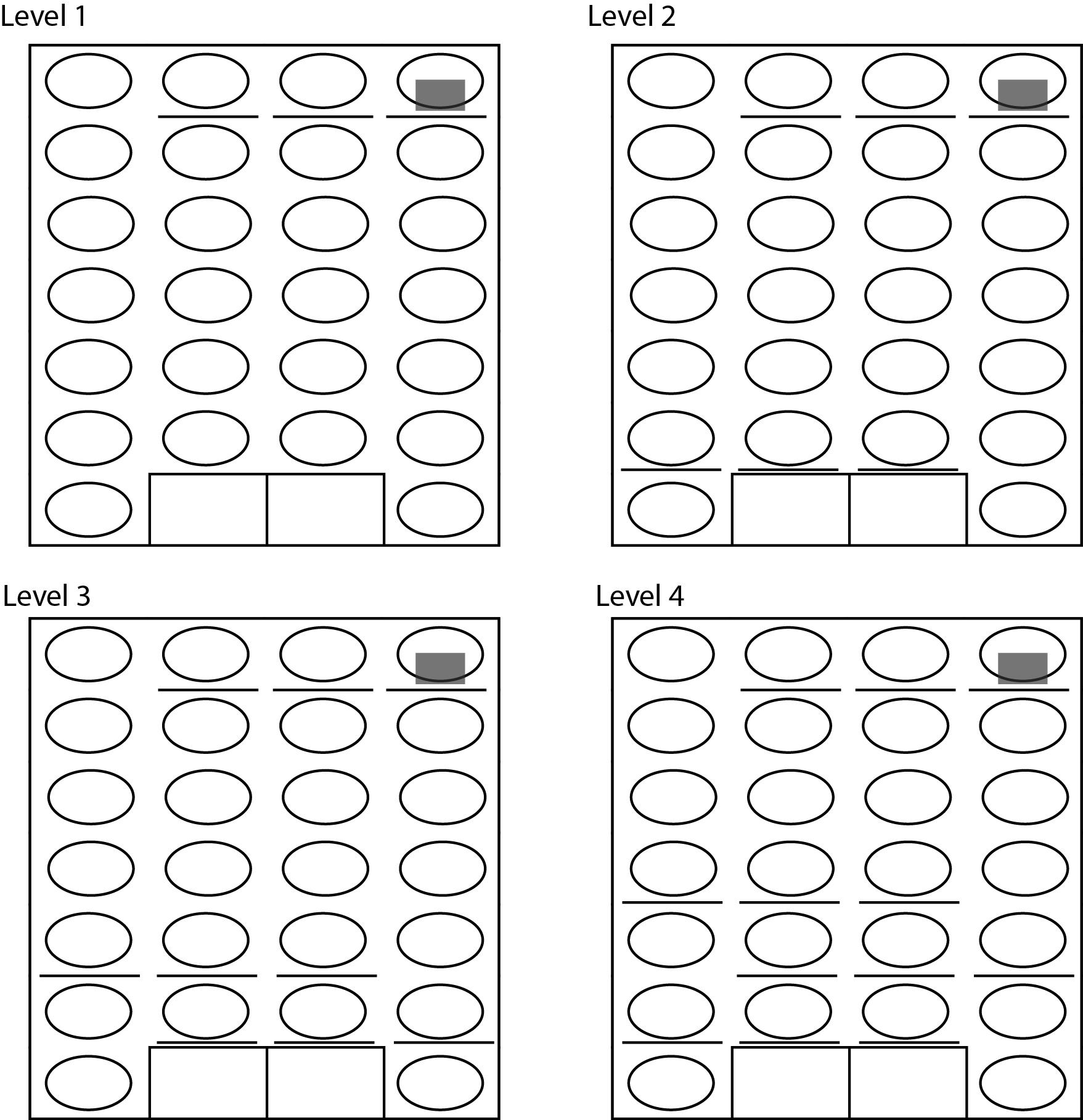

**Supplemental Figure 7.** Puzzle feeder schema. Open circles represent open areas, the grey rectangle represents the food treat, the horizontal lines represent the movable wall placement, and the rectangles represent the larger open area where the food treats can be removed at the bottom of the puzzle feeder.

**Supplemental Table 1.** Maternal virologic and PRNT comparison between ZIKV subgroups

| **Outcome** | **Comparison** | **p-value** |
| --- | --- | --- |
| Maternal plasma viremia duration | DENV_ZIKV vs. ZIKV-DAK_45g | 0.7234 |
|  | DENV_ZIKV vs. ZIKV-PR_30gd | 0.6100 |
|  | DENV_ZIKV vs. ZIKV-PR_45gd | 0.7418 |
|  | ZIKV-DAK_45g vs. ZIKV-PR_30gd | 0.9663 |
|  | ZIKV-DAK_45g vs. ZIKV-PR_45gd | 0.9670 |
|  | ZIKV-PR_30gd vs. ZIKV-PR_45gd | 0.9961 |
| Area under the curve of maternal plasma viremia | DENV_ZIKV vs. ZIKV-DAK_45g | 0.8742 |
|  | DENV_ZIKV vs. ZIKV-PR_30gd | 0.4811 |
|  | DENV_ZIKV vs. ZIKV-PR_45gd | 0.8050 |
|  | ZIKV-DAK_45g vs. ZIKV-PR_30gd | 0.8673 |
|  | ZIKV-DAK_45g vs. ZIKV-PR_45gd | 0.9266 |
|  | ZIKV-PR_30gd vs. ZIKV-PR_45gd | 0.9898 |
| Combined biopsies vRNA-positive (%) | DENV_ZIKV vs. ZIKV-DAK_45g | 0.8886 |
|  | DENV_ZIKV vs. ZIKV-PR_30gd | 0.9467 |
|  | DENV_ZIKV vs. ZIKV-PR_45gd | 0.9654 |
|  | ZIKV-DAK_45g vs. ZIKV-PR_30gd | 0.9815 |
|  | ZIKV-DAK_45g vs. ZIKV-PR_45gd | 0.7159 |
|  | ZIKV-PR_30gd vs. ZIKV-PR_45gd | 0.6356 |
| PRNT90 | DENV_ZIKV vs. ZIKV-DAK_45g | 0.7055 |
|  | DENV/ZIKV vs. ZIKV-PR 30gd | 0.3146 |
|  | DENV/ZIKV vs. ZIKV-PR 45gd | 0.3683 |
|  | ZIKV-DAK_45g vs. ZIKV-PR_30gd | 0.1649 |
|  | ZIKV-DAK_45g vs. ZIKV-PR_45gd | 0.6434 |
|  | ZIKV-PR 30gd vs. ZIKV-PR 45gd | 0.1851 |
| Placental biopsies vRNA-positive (%) | DENV_ZIKV vs. ZIKV-DAK_45g | 0.1804 |
|  | DENV/ZIKV vs. ZIKV-PR 30gd | 0.1917 |
|  | DENV/ZIKV vs. ZIKV-PR 45gd | 0.2400 |
|  | ZIKV-DAK_45g vs. ZIKV-PR_30gd | 0.8236 |
|  | ZIKV-DAK_45g vs. ZIKV-PR_45gd | 0.9110 |
|  | ZIKV-PR 30gd vs. ZIKV-PR 45gd | 1.0000 |
| Decidual biopsies vRNA-positive (%) | DENV_ZIKV vs. ZIKV-DAK_45g | 0.7184 |
|  | DENV/ZIKV vs. ZIKV-PR 30gd | 0.4480 |
|  | DENV/ZIKV vs. ZIKV-PR 45gd | 0.4485 |
|  | ZIKV-DAK_45g vs. ZIKV-PR_30gd | 0.4783 |
|  | ZIKV-DAK_45g vs. ZIKV-PR_45gd | 0.4794 |
|  | ZIKV-PR 30gd vs. ZIKV-PR 45gd | 0.8562 |
| Chorionic membrane biopsies vRNA-positive (%) | DENV_ZIKV vs. ZIKV-DAK_45g | 0.7184 |
|  | DENV/ZIKV vs. ZIKV-PR 30gd | 0.3628 |
|  | DENV/ZIKV vs. ZIKV-PR 45gd | 0.7940 |
|  | ZIKV-DAK_45g vs. ZIKV-PR_30gd | 0.5139 |
|  | ZIKV-DAK_45g vs. ZIKV-PR_45gd | 0.7048 |
|  | ZIKV-PR 30gd vs. ZIKV-PR 45gd | 0.0738 |

**Supplemental Table 2.** Longitudinal infant blood and urine ZIKV viral loads

| **Infant ID** | **Treatment group** | **Infant age (days)** | **Specimen type (blood/ urine)** | **ZIKV vRNA copies/ml** | **ZIKV vRNA load (limit of detection of 150 copies/ml)** |
| --- | --- | --- | --- | --- | --- |
| 042-504 | DENV/ZIKV | 0 | blood | 0 | Undetectable |
| 042-504 | DENV/ZIKV | 86 | blood | 0 | Undetectable |
| 042-504 | DENV/ZIKV | 188 | blood | 0 | Undetectable |
| 042-504 | DENV/ZIKV | 358 | blood | 0 | Undetectable |
| 042-502 | DENV/ZIKV | 0 | blood | 0 | Undetectable |
| 042-502 | DENV/ZIKV | 89 | blood | 0 | Undetectable |
| 042-502 | DENV/ZIKV | 188 | blood | 0 | Undetectable |
| 042-502 | DENV/ZIKV | 359 | blood | 0 | Undetectable |
| 042-501 | DENV/ZIKV | 3 | blood | 0 | Undetectable |
| 042-501 | DENV/ZIKV | 89 | blood | 0 | Undetectable |
| 042-501 | DENV/ZIKV | 181 | blood | 0 | Undetectable |
| 042-501 | DENV/ZIKV | 357 | blood | 0 | Undetectable |
| 044-504 | ZIKV-PR 45gd | 0 | blood | 0 | Undetectable |
| 044-504 | ZIKV-PR 45gd | 0 | urine | 0 | Undetectable |
| 044-504 | ZIKV-PR 45gd | 86 | blood | 0 | Undetectable |
| 044-504 | ZIKV-PR 45gd | 183 | blood | 0 | Undetectable |
| 044-504 | ZIKV-PR 45gd | 357 | blood | 0 | Undetectable |
| 044-503 | ZIKV-PR 45gd | 0 | blood | 0 | Undetectable |
| 044-503 | ZIKV-PR 45gd | 0 | urine | 0 | Undetectable |
| 044-503 | ZIKV-PR 45gd | 86 | blood | 0 | Undetectable |
| 044-503 | ZIKV-PR 45gd | 183 | blood | 0 | Undetectable |
| 044-503 | ZIKV-PR 45gd | 363 | blood | 0 | Undetectable |
| 044-502 | ZIKV-PR 45gd | 0 | urine | 0 | Undetectable |
| 044-502 | ZIKV-PR 45gd | 0 | urine | 0 | Undetectable |
| 044-502 | ZIKV-PR 45gd | 88 | blood | 0 | Undetectable |
| 044-502 | ZIKV-PR 45gd | 175 | blood | 0 | Undetectable |
| 044-502 | ZIKV-PR 45gd | 368 | blood | 0 | Undetectable |
| 044-501 | ZIKV-PR 45gd | 0 | urine | 0 | Undetectable |
| 044-501 | ZIKV-PR 45gd | 0 | blood | 0 | Undetectable |
| 044-501 | ZIKV-PR 45gd | 86 | blood | 0 | Undetectable |
| 044-501 | ZIKV-PR 45gd | 177 | blood | 0 | Undetectable |
| 044-501 | ZIKV-PR 45gd | 361 | blood | 0 | Undetectable |
| 044-507 | Control | 173 | blood | 0 | Undetectable |
| 044-507 | Control | 392 | blood | 0 | Undetectable |
| 044-505 | Control | 178 | blood | 0 | Undetectable |
| 044-506 | Control | 176 | blood | 0 | Undetectable |
| 042-507 | DENV/ZIKV | 0 | blood | 0 | Undetectable |
| 042-507 | DENV/ZIKV | 88 | blood | 0 | Undetectable |
| 042-507 | DENV/ZIKV | 176 | blood | 0 | Undetectable |
| 042-507 | DENV/ZIKV | 361 | blood | 0 | Undetectable |
| 044-508 | Control | 184 | blood | 0 | Undetectable |
| 042-505 | DENV/ZIKV | 0 | blood | 0 | Undetectable |
| 042-505 | DENV/ZIKV | 93 | blood | 0 | Undetectable |
| 042-505 | DENV/ZIKV | 176 | blood | 0 | Undetectable |
| 042-505 | DENV/ZIKV | 371 | blood | 0 | Undetectable |
| 042-508 | DENV/ZIKV | 0 | blood | 0 | Undetectable |
| 042-508 | DENV/ZIKV | 0 | urine | 0 | Undetectable |
| 042-508 | DENV/ZIKV | 92 | blood | 0 | Undetectable |
| 042-508 | DENV/ZIKV | 183 | blood | 0 | Undetectable |
| 042-508 | DENV/ZIKV | 365 | blood | 0 | Undetectable |
| 042-506 | DENV/ZIKV | 0 | blood | 0 | Undetectable |
| 042-506 | DENV/ZIKV | 90 | blood | 0 | Undetectable |
| 042-506 | DENV/ZIKV | 176 | blood | 0 | Undetectable |
| 042-506 | DENV/ZIKV | 365 | blood | 0 | Undetectable |
| 046-501 | ZIKV-DAK | 0 | blood | 0 | Undetectable |
| 046-501 | ZIKV-DAK | 7 | blood | 0 | Undetectable |
| 046-501 | ZIKV-DAK | 92 | blood | 0 | Undetectable |
| 046-501 | ZIKV-DAK | 94 | blood | 0 | Undetectable |
| 046-501 | ZIKV-DAK | 269 | blood | 0 | Undetectable |
| 046-501 | ZIKV-DAK | 372 | blood | 0 | Undetectable |
| 046-502 | ZIKV-DAK | 0 | blood | 0 | Undetectable |
| 046-502 | ZIKV-DAK | 0 | urine | 0 | Undetectable |
| 046-502 | ZIKV-DAK | 97 | blood | 0 | Undetectable |
| 046-502 | ZIKV-DAK | 267 | blood | 0 | Undetectable |
| 046-502 | ZIKV-DAK | 370 | blood | 0 | Undetectable |
| 044-509 | ZIKV-PR 45gd | 7 | blood | 0 | Undetectable |
| 044-509 | ZIKV-PR 45gd | 43 | blood | 0 | Undetectable |
| 044-509 | ZIKV-PR 45gd | 87 | blood | 0 | Undetectable |
| 044-509 | ZIKV-PR 45gd | 184 | blood | 0 | Undetectable |
| 044-509 | ZIKV-PR 45gd | 365 | blood | 0 | Undetectable |
| 044-510 | ZIKV-PR 30gd | 0 | blood | 0 | Undetectable |
| 044-510 | ZIKV-PR 30gd | 108 | blood | 0 | Undetectable |
| 044-510 | ZIKV-PR 30gd | 116 | blood | 0 | Undetectable |
| 044-510 | ZIKV-PR 30gd | 177 | blood | 0 | Undetectable |
| 044-510 | ZIKV-PR 30gd | 360 | blood | 0 | Undetectable |
| 044-511 | Control | NA | NA | NA | NA |
| 044-512 | ZIKV-PR 30gd | 0 | blood | 0 | Undetectable |
| 044-512 | ZIKV-PR 30gd | 90 | blood | 0 | Undetectable |
| 044-512 | ZIKV-PR 30gd | 182 | blood | 0 | Undetectable |
| 044-512 | ZIKV-PR 30gd | 356 | blood | 0 | Undetectable |
| 044-514 | ZIKV-PR 30gd | 0 | blood | 0 | Undetectable |
| 044-514 | ZIKV-PR 30gd | 92 | blood | 0 | Undetectable |
| 044-514 | ZIKV-PR 30gd | 177 | blood | 0 | Undetectable |
| 044-514 | ZIKV-PR 30gd | 337 | blood | 0 | Undetectable |
| 044-513 | Control | NA | NA | NA | NA |
| 044-515 | Control | 49 | blood | 0 | Undetectable |
| 044-515 | Control | 91 | blood | 0 | Undetectable |
| 044-516 | ZIKV-PR 30gd | 0 | blood | 0 | Undetectable |
| 044-516 | ZIKV-PR 30gd | 48 | blood | 0 | Undetectable |
| 044-516 | ZIKV-PR 30gd | 95 | blood | 0 | Undetectable |
| 044-516 | ZIKV-PR 30gd | 169 | blood | 0 | Undetectable |
| 044-516 | ZIKV-PR 30gd | 350 | blood | 0 | Undetectable |
| 046-505 | ZIKV-DAK | 0 | blood | 0 | Undetectable |
| 046-505 | ZIKV-DAK | 0 | urine | 0 | Undetectable |
| 046-505 | ZIKV-DAK | 49 | blood | 0 | Undetectable |
| 046-505 | ZIKV-DAK | 91 | blood | 0 | Undetectable |
| 046-505 | ZIKV-DAK | 182 | blood | 0 | Undetectable |
| 046-505 | ZIKV-DAK | 353 | blood | 0 | Undetectable |
| 044-517 | ZIKV-PR 30gd | 0 | blood | 0 | Undetectable |
| 044-517 | ZIKV-PR 30gd | 50 | blood | 0 | Undetectable |
| 044-517 | ZIKV-PR 30gd | 101 | blood | 0 | Undetectable |
| 044-517 | ZIKV-PR 30gd | 219 | blood | 109.35 | Undetectable |
| 044-517 | ZIKV-PR 30gd | 373 | blood | 0 | Undetectable |
| 044-518 | ZIKV-PR 30gd | 0 | blood | 0 | Undetectable |
| 044-518 | ZIKV-PR 30gd | 50 | blood | 0 | Undetectable |
| 044-518 | ZIKV-PR 30gd | 84 | blood | 0 | Undetectable |
| 044-518 | ZIKV-PR 30gd | 172 | blood | 0 | Undetectable |
| 044-518 | ZIKV-PR 30gd | 350 | blood | 0 | Undetectable |
| 046-506 | ZIKV-DAK | 0 | urine | 0 | Undetectable |
| 046-506 | ZIKV-DAK | 0 | blood | 0 | Undetectable |
| 046-506 | ZIKV-DAK | 48 | blood | 0 | Undetectable |
| 046-506 | ZIKV-DAK | 83 | blood | 0 | Undetectable |
| 046-506 | ZIKV-DAK | 182 | blood | 0 | Undetectable |
| 046-506 | ZIKV-DAK | 362 | blood | 0 | Undetectable |
| 044-520 | Control | NA | NA | NA | NA |
| 044-523 | Control | NA | NA | NA | NA |
| 044-522 | ZIKV-PR 30gd | 0 | blood | 0 | Undetectable |
| 044-522 | ZIKV-PR 30gd | 0 | blood | 0 | Undetectable |
| 044-522 | ZIKV-PR 30gd | 43 | blood | 0 | Undetectable |
| 044-522 | ZIKV-PR 30gd | 97 | blood | 0 | Undetectable |
| 044-522 | ZIKV-PR 30gd | 183 | blood | 0 | Undetectable |
| 044-522 | ZIKV-PR 30gd | 361 | blood | 0 | Undetectable |
| 044-524 | Control | NA | NA | NA | NA |
| 044-525 | Control | NA | NA | NA | NA |
| 044-526 | ZIKV-PR 45gd | 0 | blood | 0 | Undetectable |
| 044-526 | ZIKV-PR 45gd | 41 | blood | 0 | Undetectable |
| 044-526 | ZIKV-PR 45gd | 91 | blood | 0 | Undetectable |
| 044-526 | ZIKV-PR 45gd | 178 | blood | 0 | Undetectable |
| 044-526 | ZIKV-PR 45gd | 363 | blood | 0 | Undetectable |
| 044-527 | ZIKV-PR 45gd | 7 | blood | 0 | Undetectable |
| 044-527 | ZIKV-PR 45gd | 48 | blood | 0 | Undetectable |
| 044-527 | ZIKV-PR 45gd | 88 | blood | 0 | Undetectable |
| 044-527 | ZIKV-PR 45gd | 183 | blood | 0 | Undetectable |
| 044-527 | ZIKV-PR 45gd | 363 | blood | 0 | Undetectable |
| 044-528 | Control | NA | NA | NA | NA |
| 044-530 | ZIKV-PR 45gd | 0 | blood | 0 | Undetectable |
| 044-530 | ZIKV-PR 45gd | 43 | blood | 0 | Undetectable |
| 044-530 | ZIKV-PR 45gd | 94 | blood | 0 | Undetectable |
| 044-530 | ZIKV-PR 45gd | 176 | blood | 0 | Undetectable |
| 044-530 | ZIKV-PR 45gd | 361 | blood | 0 | Undetectable |
| 044-531 | ZIKV-PR 30gd | 6 | blood | 0 | Undetectable |
| 044-531 | ZIKV-PR 30gd | 47 | blood | 0 | Undetectable |
| 044-531 | ZIKV-PR 30gd | 93 | blood | 0 | Undetectable |
| 044-531 | ZIKV-PR 30gd | 173 | blood | 0 | Undetectable |
| 044-531 | ZIKV-PR 30gd | 359 | blood | 0 | Undetectable |
| 044-532 | ZIKV-PR 30gd | 0 | blood | 0 | Undetectable |
| 044-532 | ZIKV-PR 30gd | 46 | blood | 0 | Undetectable |
| 044-532 | ZIKV-PR 30gd | 91 | blood | 0 | Undetectable |
| 044-532 | ZIKV-PR 30gd | 173 | blood | 0 | Undetectable |
| 044-533 | ZIKV-PR 45gd | 0 | blood | 0 | Undetectable |
| 044-533 | ZIKV-PR 45gd | 42 | blood | 0 | Undetectable |
| 044-533 | ZIKV-PR 45gd | 109 | blood | 0 | Undetectable |
| 044-533 | ZIKV-PR 45gd | 176 | blood | 0 | Undetectable |
| 044-533 | ZIKV-PR 45gd | 358 | blood | 0 | Undetectable |

**Supplemental Table 3**. ZIKV-exposed infant IgM testing

| **Infant ID** | **Age (days)** | **ZIKV-specific IgM testing** |
| --- | --- | --- |
| 042-505 | 0 | Negative |
| 042-506 | 0 | Negative |
| 042-507 | 0 | Negative |
| 042-508 | 0 | Negative |
| 044-501 | 0 | Negative |
| 044-502 | 0 | Negative |
| 044-503 | 0 | Negative |
| 044-504 | 0 | Negative |
| 044-509 | 7 | Negative |
| 044-510 | 0 | Negative |
| 044-512 | 0 | Negative |
| 044-514 | 0 | Negative |
| 044-516 | 0 | Negative |
| 044-517 | 0 | Negative |
| 044-522 | 0 | Negative |
| 044-526 | 0 | Negative |
| 044-527 | 7 | Negative |
| 044-530 | 0 | Negative |
| 044-531 | 8 | Negative |
| 044-532 | 0 | Negative |
| 044-533 | 0 | Negative |
| 046-501 | 0 | Negative |
| 046-502 | 0 | Negative |
| 046-505 | 0 | Negative |
| 046-506 | 0 | Negative |

Four ZIKV-exposed infants did not have serum or plasma available from the first week of life and could not be tested.

**Supplemental Table 4.** Infant demographics

| **Demographic parameter** | **Treatment group** | **Mean (95% CI*)** | **p-value (All ZIKV vs Control)** | **Number of infants included in specific analysis** |
| --- | --- | --- | --- | --- |
| Gestational age at maternal inoculation (days) | DENV/ZIKV | 47 (43-51) |  | 7 |
|  | ZIKV-DAKAR 45gd | 45 (39-50) |  | 4 |
|  | ZIKV-PR 30gd | 29 (26-33) |  | 9 |
|  | ZIKV-PR 45gd | 45 (42-49) |  | 9 |
|  | Controls | 40 (37-43) | 0.7683 | 12 |
|  | All ZIKV | 41 (38-44) |  | 29 |
| Delivery method by Cesarean section (%) | DENV/ZIKV | 100 (65-100) |  | 7 |
|  | ZIKV-DAKAR 45gd | 100 (51-100%) |  | 4 |
|  | ZIKV-PR 30gd | 89 ( 57-98) |  | 9 |
|  | ZIKV-PR 45gd | 89 (57-98) |  | 9 |
|  | Controls | 92 (65-99%) | 0.9999 | 12 |
|  | All ZIKV | 93 (78-98) |  | 29 |
| Male sex (%) | DENV/ZIKV | 29 (9-64) |  | 7 |
|  | ZIKV-DAKAR 45gd | 25 (5-70) |  | 4 |
|  | ZIKV-PR 30gd | 78 (45-94) |  | 9 |
|  | ZIKV-PR 45gd | 78 (45-04) |  | 9 |
|  | Controls | 0 (0-25) | 0.0014 | 12 |
|  | All ZIKV | 59 (40-74) |  | 29 |
| Housing status with dam at 12 months (%) | DENV/ZIKV | 86 (49-97) |  | 7 |
|  | ZIKV-DAKAR 45gd | 75 (30-95) |  | 4 |
|  | ZIKV-PR 30gd | 100 (70-100) |  | 9 |
|  | ZIKV-PR 45gd | 89 (57-98) |  | 9 |
|  | Controls | 75 (47-91) | 0.3344 | 12 |
|  | All ZIKV | 90 (74-96) |  | 29 |
| Age at 3 month timepoint eye exam (days) | DENV/ZIKV | 92 (88-95) |  | 7 |
|  | ZIKV-DAKAR 45gd | 90 (85-95) |  | 4 |
|  | ZIKV-PR 30gd | 95 (91-98) |  | 9 |
|  | ZIKV-PR 45gd | 93 (90-97) |  | 9 |
|  | Controls | 91 (88-94) | 0.2555 | 11 |
|  | All ZIKV | 93 (91-95) |  | 29 |
| Age at 12 month timepoint eye exam (days) | DENV/ZIKV | 361 (354-368) |  | 7 |
|  | ZIKV-DAKAR 45gd | 368 (359-377) |  | 4 |
|  | ZIKV-PR 30gd | 356 (350-362) |  | 9 |
|  | ZIKV-PR 45gd | 360 (354-366) |  | 9 |
|  | Controls | 362 (357-367) | 0.5876 | 12 |
|  | All ZIKV | 360 (357-364) |  | 29 |
| Age at 1 month timepoint hearing exam (days) | DENV/ZIKV | Not done |  | 0 |
|  | ZIKV-DAKAR 45gd | 48 (43-53) |  | 2 |
|  | ZIKV-PR 30gd | 48 (45-51) |  | 6 |
|  | ZIKV-PR 45gd | 42 (38-45) |  | 5 |
|  | Controls | 44 (41-47) | 0.3917 | 6 |
|  | All ZIKV | 46 (43-48) |  | 13 |
| Age at 3 month timepoint hearing exam (days) | DENV/ZIKV | 90 (85-95) |  | 7 |
|  | ZIKV-DAKAR 45gd | 89 (83-95) |  | 4 |
|  | ZIKV-PR 30gd | 94 (89-98) |  | 8 |
|  | ZIKV-PR 45gd | 91 (87-95) |  | 9 |
|  | Controls | 92 (88-95) | 0.8765 | 12 |
|  | All ZIKV | 91 (89-93) |  | 28 |
| Age at 6 month timepoint hearing exam (days) | DENV/ZIKV | 181 (175-188) |  | 7 |
|  | ZIKV-DAKAR 45gd | 182 (170-194) |  | 2 |
|  | ZIKV-PR 30gd | 183 (177-189) |  | 9 |
|  | ZIKV-PR 45gd | 179 (174-185) |  | 9 |
|  | Controls | 179 (174-184) | 0.3555 | 12 |
|  | All ZIKV | 181 (178-185) |  | 27 |
| Age at 12 month timepoint hearing exam (days) | DENV/ZIKV | 362 (355-368) |  | 7 |
|  | ZIKV-DAKAR 45gd | 364 (356-373) |  | 4 |
|  | ZIKV-PR 30gd | 356 (351-362) |  | 9 |
|  | ZIKV-PR 45gd | 361 (355-367) |  | 9 |
|  | Controls | 362 (357-367) | 0.4510 | 12 |
|  | All ZIKV | 360 (357-363) |  | 29 |
| Age at mother-infant dyad testing (days) | DENV/ZIKV | 364 (351-377) |  | 6 |
|  | ZIKV-DAKAR 45gd | 367 (349-385) |  | 3 |
|  | ZIKV-PR 30gd | 349 (338-359) |  | 9 |
|  | ZIKV-PR 45gd | 357 (346-368) |  | 8 |
|  | Controls | 356 (345-366) | 0.8462 | 9 |
|  | All ZIKV | 357 (350-363) |  | 26 |
| Age at first day of behavioral testing (days) | DENV/ZIKV | 363 (356-371) |  | 7 |
|  | ZIKV-DAKAR 45gd | 364 (355-374) |  | 4 |
|  | ZIKV-PR 30gd | 358 (352-364) |  | 9 |
|  | ZIKV-PR 45gd | 357 (351-363) |  | 9 |
|  | Controls | 361 (355-367) | 0.7079 | 11 |
|  | All ZIKV | 360 (356-363) |  | 29 |

*Confidence interval

^Missing values are not included in calculations

**Supplemental Table 5.** Infant head circumference and weight gain patterns

| **Timepoint comparisons** | **Outcome** | **Age (months)** | **Group** | **Mean** | **95% CI** | **p-value** |
| --- | --- | --- | --- | --- | --- | --- |
|  | Head circumference (cm) | 0 | Control | 19.7 | 19.3-20 | 0.4965 |
|  |  |  | ZIKV | 19.5 | 19.3-19.7 |  |
|  |  | 12 | Control | 24.3 | 24-24.6 | 0.1049 |
|  |  |  | ZIKV | 24.6 | 24.4-24.8 |  |
|  | Weight (kg) | 0 | Control | 0.5 | 0.4-0.6 | 0.8365 |
|  |  |  | ZIKV | 0.5 | 0.5-0.6 |  |
|  |  | 12 | Control | 2.1 | 2-2.2 | 0.0121 |
|  |  |  | ZIKV | 2.2 | 2.2-2.3 |  |
| **Trajectory comparisons** | **Outcome** | **Age (months)** | **Group** | **Slope (mm/month)** | **95% CI** | **p-value** |
|  | Head circumference | 0 to 12 | Control | 3.4 | 2.89 - 3.82 | 0.2162 |
|  |  |  | ZIKV | 3.7 | 3.39 - 4.04 |  |
|  | **Outcome** | **Age (months)** | **Group** | **Slope (g/month)** | **95% CI** | **p-value** |
|  | Weight | 0 to 12 | Control | 137.3 | 130.55 - 144.01 | 0.0317 |
|  |  |  | ZIKV | 146.1 | 141.71 - 150.47 |  |

**Supplemental Table 6.** Infant developmental testing statistical comparisons

| Outcome | Group | N | Median | IQR (Interquartile Range) | p-value |
| --- | --- | --- | --- | --- | --- |
| Mother - infant home cage observations | | | | | |
| Mutual ventral contact | ZIKV | 26 | 36.3 | 17.5 - 50 | 0.0127 |
| Mutual ventral contact | Control | 9 | 0.0 | 0 - 4 |  |
| Together | ZIKV | 26 | 74.8 | 58.3 - 90.4 | 0.0149 |
| Together | Control | 9 | 50.4 | 17.9 - 67.5 |  |
| Nipple contact | ZIKV | 26 | 33.3 | 0.9 - 49.4 | 0.0093 |
| Nipple contact | Control | 9 | 0.0 | 0 - 1.2 |  |
| Locomotion | ZIKV | 26 | 10.6 | 8.2 - 27.7 | 0.3751 |
| Locomotion | Control | 9 | 18.1 | 13.4 - 19.2 |  |
| Puzzle Feeder | | | | | |
| Number of levels completed, day 1 | ZIKV | 26 | 1 | 0.0 - 1.0 | 0.3953 |
|  | Control | 8 | 1 | 0.8 - 1.2 |  |
| Number of levels completed, day 2 | ZIKV | 26 | 1 | 0.0 - 1.0 | 0.1238 |
|  | Control | 8 | 1 | 1.0 - 1.2 |  |
| Number of levels completed, day 3 | ZIKV | 26 | 1 | 0.0 - 1.0 | 0.9999 |
|  | Control | 7 | 1 | 0.5 - 1.5 |  |
| Time to complete Level 1 (sec), day 1 | ZIKV | 25 | 830 | 352.0 - 1775.0 | 0.0389 |
|  | Control | 7 | 442 | 135.0 - 599.5 |  |
| Time to complete Level 1 (sec), day 2 | ZIKV | 25 | 398 | 195.0 - 550.0 | 0.5136 |
|  | Control | 8 | 270 | 194.8 - 362.5 |  |
| Time to complete Level 1 (sec), day 3 | ZIKV | 25 | 578 | 230.0 - 1785.0 | 0.1157 |
|  | Control | 7 | 233 | 201.5 - 535.5 |  |
| Number of attempts to complete Level 1, day 1 | ZIKV | 26 | 10 | 7.2 - 15.8 | 0.5802 |
|  | Control | 8 | 11 | 6.8 - 13.8 |  |
| Number of attempts to complete Level 1, day 2 | ZIKV | 26 | 10.5 | 4.2 - 13.8 | 0.4891 |
|  | Control | 8 | 8 | 6.5 - 12.0 |  |
| Number of attempts to complete Level 1, day 3 | ZIKV | 26 | 10 | 6.2 - 16.0 | 0.3626 |
|  | Control | 7 | 9 | 8.5 - 10.0 |  |
| Number of attempts to complete Level 2, day 1 | ZIKV | 15 | 12 | 9.0 - 18.0 | 0.3094 |
|  | Control | 6 | 9 | 4.5 - 15.0 |  |
| Number of attempts to complete Level 2, day 2 | ZIKV | 22 | 10.5 | 5.2 - 15.0 | 0.4073 |
|  | Control | 8 | 8.5 | 5.2 - 14.0 |  |
| Number of attempts to complete Level 2, day 3 | ZIKV | 18 | 6.5 | 3.2 - 13.0 | 0.9977 |
|  | Control | 7 | 7 | 5.5 - 11.5 |  |
| PVC Pipe Test | | | | | |
| Oral exploration percent of time, day 2 | ZIKV | 24 | 23.1355 | 7.9 - 29.2 | 0.0681 |
|  | Control | 9 | 11.7365 | 5.0 - 15.8 |  |
| Oral exploration percent of time, day 3 | ZIKV | 24 | 15.86275 | 7.6 - 30.8 | 0.1071 |
|  | Control | 8 | 9.724765 | 5.8 - 14.5 |  |
| Finger in frosting percent of time , day 2 | ZIKV | 25 | 23.0777 | 19.4 - 29.8 | 0.8588 |
|  | Control | 9 | 25.192 | 8.0 - 29.0 |  |
| Finger in frosting percent of time , day 3 | ZIKV | 24 | 20.47565 | 14.4 - 29.0 | 0.9218 |
|  | Control | 8 | 23.856 | 10.8 - 26.7 |  |
| Finger isolation percent of time, day 2 | ZIKV | 24 | 12.17595 | 5.6 - 19.9 | 0.1785 |
|  | Control | 9 | 4.31277 | 2.5 - 17.9 |  |
| Finger isolation percent of time, day3 | ZIKV | 24 | 7.117885 | 4.4 - 14.1 | 0.1536 |
|  | Control | 8 | 10.38437 | 6.2 - 24.7 |  |
| Tactile Response | | | | | |
|  | | N | Approach n (%) | Did not approach n (%) | p-value |
| Infant approach feather, day 1 | ZIKV | 26 | 23 (88%) | 3 (12%) | 0.0085 |
|  | Control | 8 | 3 (38%) | 5 (62%) |  |
| Infant approach feather, day 2 | ZIKV | 26 | 21 (81%) | 5 (19%) | 0.1648 |
|  | Control | 8 | 4 (50%) | 4 (50%) |  |
| Infant approach feather, day 3 | ZIKV | 26 | 22 (85%) | 4 (15%) | 1 |
|  | Control | 6 | 5 (83%) | 1 (17%) |  |
| Infant approach cottonball, day 1 | ZIKV | 26 | 24 (92%) | 2 (8%) | 0.0035 |
|  | Control | 8 | 3 (38%) | 5 (62%) |  |
| Infant approach cottonball, day 2 | ZIKV | 26 | 21 (81%) | 5 (19%) | 0.3551 |
|  | Control | 8 | 5 (62%) | 3 (38%) |  |
| Infant approach cottonball, day 3 | ZIKV | 26 | 22 (85%) | 4 (15%) | 1 |
|  | Control | 6 | 5 (83%) | 1 (17%) |  |
| Infant approach brush, day 1 | ZIKV | 26 | 23 (88%) | 3 (12%) | 0.0374 |
|  | Control | 8 | 4 (50%) | 4 (50%) |  |
| Infant approach brush, day 2 | ZIKV | 26 | 20 (77%) | 6 (23%) | 0.1949 |
|  | Control | 8 | 4 (50%) | 4 (50%) |  |
| Infant approach brush, day 3 | ZIKV | 26 | 20 (77%) | 6 (23%) | 1 |
|  | Control | 6 | 5 (83%) | 1 (17%) |  |

**Supplemental Table 7.** Ophthalmic Exam Summary

| **Infant ID** | **Treatment Group** | **Ophthalmic Exam Findings** | | |
| --- | --- | --- | --- | --- |
|  |  | **4-6 weeks** | **3 months** | **12 months** |
| 042-501 | DENV/ZIKV | X | No abnormalities | No abnormalities |
| 042-502 | DENV/ZIKV | X | No abnormalities | No abnormalities |
| 042-504 | DENV/ZIKV | X | No abnormalities | Iris nodule at 4:00 in the right eye.  Iris nodules at 4:00, 8:00, and 11:00 in the left eye. |
| 042-505 | DENV/ZIKV | X | No abnormalities | No abnormalities |
| 042-506 | DENV/ZIKV | X | No abnormalities | No abnormalities |
| 042-507 | DENV/ZIKV | X | No abnormalities | No abnormalities |
| 042-508 | DENV/ZIKV | X | No abnormalities | No abnormalities |
| 044-501 | ZIKV-PR 45gd | X | No abnormalities | No abnormalities |
| 044-502 | ZIKV-PR 45gd | X | Remnant of hyaloid system in the left eye. | No abnormalities |
| 044-503 | ZIKV-PR 45gd | X | Iris nodule at 4:30 in the right eye.  Iris nodules at 2:30, 7:30, and 8:00 in the left eye. | Iris nodule at 6:00 in the right eye.  Iris nodules at 3:00, 7:30, and 8:30 in the left eye. |
| 044-504 | ZIKV-PR 45gd | X | No abnormalities | No abnormalities |
| 044-505 | Control | X | No abnormalities | No abnormalities |
| 044-506 | Control | X | No abnormalities | No abnormalities |
| 044-507 | Control | X | No abnormalities | No abnormalities |
| 044-508 | Control | X | No abnormalities | No abnormalities |
| 044-509 | ZIKV-PR 30gd | X | No abnormalities | No abnormalities |
| 044-510 | ZIKV-PR 30gd | X | No abnormalities | No abnormalities |
| 044-511 | Control | X | A small puntate pigmentary defect located in the far temporal macula on the right | No abnormalities |
| 044-512 | ZIKV-PR 30gd | X | No abnormalities | No abnormalities |
| 044-513 | Control | X | No abnormalities | No abnormalities |
| 044-514 | ZIKV-PR 30gd | X | No abnormalities | No abnormalities |
| 044-515 | Control | No abnormalities | No abnormalities | No abnormalities |
| 044-516 | ZIKV-PR 30gd | No abnormalities | No abnormalities | No abnormalities |
| 044-517 | ZIKV-PR 30gd | No abnormalities | No abnormalities | No abnormalities |
| 044-518 | ZIKV-PR 30gd | Persistent tunica vasculosa bilaterally | Persistent tunica vasculosa on the left, resolved on the right | No abnormalities |
| 044-520 | Control | No abnormalities | No abnormalities | No abnormalities |
| 044-522 | ZIKV-PR 30gd | No abnormalities | No abnormalities | No abnormalities |
| 044-523 | Control | No abnormalities | No abnormalities | No abnormalities |
| 044-524 | Control | No abnormalities | No abnormalities | No abnormalities |
| 044-525 | Control | No abnormalities | X | No abnormalities |
| 044-526 | ZIKV-PR 45gd | No abnormalities | No abnormalities | No abnormalities |
| 044-527 | ZIKV-PR 45gd | No abnormalities | No abnormalities | No abnormalities |
| 044-528 | Control | No abnormalities | No abnormalities | No abnormalities |
| 044-530 | ZIKV-PR 45gd | Persistent tunica vasculosa on the left, none on the right | No abnormalities | No abnormalities |
| 044-531 | ZIKV-PR 30gd | No abnormalities | No abnormalities | No abnormalities |
| 044-532 | ZIKV-PR 30gd | No abnormalities | No abnormalities | No abnormalities |
| 044-533 | ZIKV-PR 45gd | Persistent tunica vasculosa bilaterally | Prominent Bergmeister papilla bilaterally | No abnormalities |
| 046-501 | ZIKV-DAK | X | No abnormalities | No abnormalities |
| 046-502 | ZIKV-DAK | X | No abnormalities | No abnormalities |
| 046-505 | ZIKV-DAK | No abnormalities | No abnormalities | No abnormalities |
| 046-506 | ZIKV-DAK | Persistent tunica vasculosa bilaterally | No abnormalities | No abnormalities |

X, exam not done at that time point

**Supplemental Table 8.** Retinal layer thickness comparisons

| **Retinal layer** | **Age (months)** | **Group** | **Adjusted Mean** | **95%CI** | **p-value** |
| --- | --- | --- | --- | --- | --- |
| Ganglion Cell Layer | 1 | Control | 0.025 | 0.023-0.027 | 0.1889 |
|  | 1 | ZIKV | 0.026 | 0.025-0.027 |  |
|  | 3 | Control | 0.024 | 0.022-0.025 | 0.0989 |
|  | 3 | ZIKV | 0.025 | 0.024-0.026 |  |
|  | 12 | Control | 0.025 | 0.024-0.027 | 0.1416 |
|  | 12 | ZIKV | 0.024 | 0.023-0.025 |  |
| Inner Nuclear Layer | 1 | Control | 0.035 | 0.032-0.037 | 0.9186 |
|  | 1 | ZIKV | 0.034 | 0.033-0.036 |  |
|  | 3 | Control | 0.033 | 0.031-0.034 | 0.8785 |
|  | 3 | ZIKV | 0.032 | 0.031-0.033 |  |
|  | 12 | Control | 0.033 | 0.031-0.035 | 0.6458 |
|  | 12 | ZIKV | 0.032 | 0.031-0.033 |  |
| Inner Plexiform Layer | 1 | Control | 0.039 | 0.036-0.041 | 0.4423 |
|  | 1 | ZIKV | 0.038 | 0.036-0.039 |  |
|  | 3 | Control | 0.038 | 0.036-0.04 | 0.1753 |
|  | 3 | ZIKV | 0.036 | 0.035-0.037 |  |
|  | 12 | Control | 0.039 | 0.037-0.04 | 0.4405 |
|  | 12 | ZIKV | 0.038 | 0.037-0.039 |  |
| Outer Nuclear Layer | 1 | Control | 0.066 | 0.062-0.07 | 0.9541 |
|  | 1 | ZIKV | 0.066 | 0.063-0.069 |  |
|  | 3 | Control | 0.072 | 0.068-0.075 | 0.9716 |
|  | 3 | ZIKV | 0.072 | 0.07-0.074 |  |
|  | 12 | Control | 0.073 | 0.07-0.076 | 0.1634 |
|  | 12 | ZIKV | 0.076 | 0.074-0.078 |  |
| Outer Plexiform Layer | 1 | Control | 0.024 | 0.022-0.026 | 0.8793 |
|  | 1 | ZIKV | 0.024 | 0.022-0.025 |  |
|  | 3 | Control | 0.025 | 0.024-0.026 | 0.0787 |
|  | 3 | ZIKV | 0.023 | 0.023-0.024 |  |
|  | 12 | Control | 0.024 | 0.023-0.026 | 0.0767 |
|  | 12 | ZIKV | 0.023 | 0.022-0.024 |  |
| Photoreceptor Inner Segment | 1 | Control | 0.029 | 0.027-0.03 | 0.5302 |
|  | 1 | ZIKV | 0.028 | 0.027-0.029 |  |
|  | 3 | Control | 0.03 | 0.028-0.031 | 0.5929 |
|  | 3 | ZIKV | 0.03 | 0.029-0.031 |  |
|  | 12 | Control | 0.031 | 0.03-0.032 | 0.4956 |
|  | 12 | ZIKV | 0.03 | 0.03-0.031 |  |
| Photoreceptor Outer Segment | 1 | Control | 0.044 | 0.041-0.048 | 0.4487 |
|  | 1 | ZIKV | 0.043 | 0.04-0.045 |  |
|  | 3 | Control | 0.046 | 0.044-0.049 | 0.3126 |
|  | 3 | ZIKV | 0.045 | 0.043-0.046 |  |
|  | 12 | Control | 0.049 | 0.047-0.052 | 0.2772 |
|  | 12 | ZIKV | 0.048 | 0.046-0.049 |  |
| Retinal Nerve Fiber Layer | 1 | Control | 0.018 | 0.017-0.02 | 0.3988 |
|  | 1 | ZIKV | 0.019 | 0.018-0.02 |  |
|  | 3 | Control | 0.018 | 0.018-0.019 | 0.3349 |
|  | 3 | ZIKV | 0.018 | 0.017-0.019 |  |
|  | 12 | Control | 0.02 | 0.019-0.021 | 0.4757 |
|  | 12 | ZIKV | 0.02 | 0.019-0.02 |  |
| Total Retinal Thickness | 1 | Control | 0.025 | 0.023-0.027 | 0.1889 |
|  | 1 | ZIKV | 0.026 | 0.025-0.027 |  |
|  | 3 | Control | 0.024 | 0.022-0.025 | 0.0989 |
|  | 3 | ZIKV | 0.025 | 0.024-0.026 |  |
|  | 12 | Control | 0.025 | 0.024-0.027 | 0.1416 |
|  | 12 | ZIKV | 0.024 | 0.023-0.025 |  |

**Supplemental Table 9.** Longitudinal auditory responses of infants with abnormal 12 month Wave IV responses.

| **Treatment group** | **Infant ID** | **Age (months)** | **Number of ears with no Wave IV response** | | | | | | | | | |
| --- | --- | --- | --- | --- | --- | --- | --- | --- | --- | --- | --- | --- |
|  |  |  | **1000 Hz stimulus** | | | **500 Hz stimulus** | | | **Click stimulus** | | | |
|  |  |  | **40 dB** | **60 dB** | **80 dB** | **40 dB** | **60 dB** | **80 dB** | **20 dB** | **30 dB** | **50 dB** | **70 dB** |
| DENV/ZIKV | 042-502 | 1 | x | x | x | x | x | x | x | x | x | x |
| DENV/ZIKV | 042-502 | 3 | 0 | 0 | 0 | x | x | x | x | 0 | 0 | 0 |
| DENV/ZIKV | 042-502 | 6 | x | x | x | x | x | x | x | 0 | 0 | 0 |
| DENV/ZIKV | 042-502 | 12 | **1** | 0 | 0 | x | x | x | x | 0 | 0 | 0 |
| ZIKV PR GD45 | 044-502 | 1 | x | x | x | x | x | x | x | x | x | x |
| ZIKV PR GD45 | 044-502 | 3 | **2** | **1** | 0 | x | x | x | x | **1** | 0 | 0 |
| ZIKV PR GD45 | 044-502 | 6 | 0 | 0 | 0 | x | x | x | x | 0 | 0 | 0 |
| ZIKV PR GD45 | 044-502 | 12 | 0 | 0 | 0 | **2** | 0 | 0 | 0 | 0 | 0 | 0 |
| Control | 044-523 | 1 | **2** | 0 | 0 | x | x | x | **2** | 0 | 0 | 0 |
| Control | 044-523 | 3 | x | x | x | x | x | x | 0 | 0 | 0 | 0 |
| Control | 044-523 | 6 | 0 | 0 | 0 | 0 | 0 | 0 | 0 | 0 | 0 | 0 |
| Control | 044-523 | 12 | **1** | 0 | 0 | **2** | 0 | 0 | 0 | 0 | 0 | 0 |
| ZIKV PR GD30 | 044-510 | 1 | x | x | x | x | x | x | x | x | x | x |
| ZIKV PR GD30 | 044-510 | 3 | **1** | 0 | 0 | x | x | x | x | 0 | 0 | 0 |
| ZIKV PR GD30 | 044-510 | 6 | 0 | 0 | 0 | 0 | 0 | 0 | 0 | 0 | 0 | 0 |
| ZIKV PR GD30 | 044-510 | 12 | **1** | 0 | 0 | 0 | 0 | 0 | 0 | 0 | 0 | 0 |
| ZIKV PR GD30 | 044-514 | 1 | x | x | x | x | x | x | x | x | x | x |
| ZIKV PR GD30 | 044-514 | 3 | x | x | x | x | x | x | 0 | 0 | 0 | 0 |
| ZIKV PR GD30 | 044-514 | 6 | 0-L, X-R | 0-L, X-R | 0-L, X-R | 0-L, X-R | 0-L, X-R | 0-L, X-R | **1** | **1** | **1** | 0 |
| ZIKV PR GD30 | 044-514 | 12 | 0 | 0 | 0 | **2** | 0 | 0 | 0 | 0 | 0 | 0 |
| ZIKV PR GD30 | 044-518 | 1 | **1** | 0 | 0 | **1** | **1** | 0 | **1** | 0 | 0 | 0 |
| ZIKV PR GD30 | 044-518 | 3 | 0 | 0 | 0 | 0 | 0 | 0 | 0 | 0 | 0 | 0 |
| ZIKV PR GD30 | 044-518 | 6 | 0 | 0 | 0 | 0 | 0 | 0 | 0 | 0 | 0 | 0 |
| ZIKV PR GD30 | 044-518 | 12 | 0 | 0 | 0 | **1** | 0 | 0 | 0 | 0 | 0 | 0 |

X = test was not done because of sedation problems, COVID cancellations, or the time point or specific testing variable was not implemented until later in the study. One infant had testing performed on the left side with no waveform IV abnormalities noted (0-L) but could not have the right ear examined because of problems waking up from sedation (X-R).

**Supplemental Table 10.** Dam demographics, inoculation schema, and associated offspring

| Dam ID | Inoculation type | Gestational age at inoculation | DENV infection prior to pregnancy | Treatment group name | Infant ID |
| --- | --- | --- | --- | --- | --- |
| 042-101 | ZIKV-PR | 48 | yes | DENV/ZIKV | 042-501 |
| 042-102 | ZIKV-PR | 46 | yes | DENV/ZIKV | 042-502 |
| 042-104 | ZIKV-PR | 46 | yes | DENV/ZIKV | 042-504 |
| 042-105 | ZIKV-PR | 49 | yes | DENV/ZIKV | 042-505 |
| 042-106 | ZIKV-PR | 46 | yes | DENV/ZIKV | 042-506 |
| 042-107 | ZIKV-PR | 48 | yes | DENV/ZIKV | 042-507 |
| 042-108 | ZIKV-PR | 49 | yes | DENV/ZIKV | 042-508 |
| 044-101 | ZIKV-PR | 45 | - | ZIKV-PR 45gd* | 044-501 |
| 044-102 | ZIKV-PR | 45 | - | ZIKV-PR 45gd | 044-502 |
| 044-103 | ZIKV-PR | 44 | - | ZIKV-PR 45gd | 044-503 |
| 044-104 | ZIKV-PR | 45 | - | ZIKV-PR 45gd | 044-504 |
| 044-105 | PBS inoculation | 42 | - | Control | 044-505 |
| 044-106 | PBS inoculation | 48 | - | Control | 044-506 |
| 044-107 | PBS inoculation | 48 | - | Control | 044-507 |
| 044-108 | PBS inoculation | 48 | - | Control | 044-508 |
| 044-109 | ZIKV-PR | 48 | - | ZIKV-PR 45gd | 044-509 |
| 044-110 | ZIKV-PR | 28 | - | ZIKV-PR 30gd | 044-510 |
| 044-111 | PBS inoculation | 26 | - | Control | 044-511 |
| 044-112 | ZIKV-PR | 30 | - | ZIKV-PR 30gd | 044-512 |
| 044-113 | PBS inoculation | 26 | - | Control | 044-513 |
| 044-114 | ZIKV-PR | 25 | - | ZIKV-PR 30gd | 044-514 |
| 044-115 | PBS inoculation | 30 | - | Control | 044-515 |
| 044-116 | ZIKV-PR | 26 | - | ZIKV-PR 30gd | 044-516 |
| 044-117 | ZIKV-PR | 30 | - | ZIKV-PR 30gd | 044-517 |
| 044-118 | ZIKV-PR | 30 | - | ZIKV-PR 30gd | 044-518 |
| 044-120 | PBS inoculation | 30 | - | Control | 044-520 |
| 044-122 | ZIKV-PR | 33 | - | ZIKV-PR 30gd | 044-522 |
| 020-101 | PBS inoculation | 45 | - | Control | 044-523 |
| 044-124 | PBS inoculation | 45 | - | Control | 044-524 |
| 044-125 | PBS inoculation | 45 | - | Control | 044-525 |
| 044-126 | ZIKV-PR | 44 | - | ZIKV-PR 45gd | 044-526 |
| 044-127 | ZIKV-PR | 45 | - | ZIKV-PR 45gd | 044-527 |
| 044-128 | PBS inoculation | 45 | - | Control | 044-528 |
| 044-130 | ZIKV-PR | 46 | - | ZIKV-PR 45gd | 044-530 |
| 044-131 | ZIKV-PR | 29 | - | ZIKV-PR 30gd | 044-531 |
| 044-132 | ZIKV-PR | 31 | - | ZIKV-PR 30gd | 044-532 |
| 044-133 | ZIKV-PR | 45 | - | ZIKV-PR 45gd | 044-533 |
| 046-101 | ZIKV-DAK | 48 | - | ZIKV-DAK | 046-501 |
| 046-102 | ZIKV-DAK | 46 | - | ZIKV-DAK | 046-502 |
| 046-105 | ZIKV-DAK | 43 | - | ZIKV-DAK | 046-505 |
| 046-106 | ZIKV-DAK | 42 | - | ZIKV-DAK | 046-506 |

*Gestational day (gd).

**Supplemental Table 11.** Infant and delivery demographics

| Infant ID | Delivery method (Natural or Cesarean section) | Infant sex | Gestational age at delivery | Birth weight (kg) | Housing status (Dam or peer) |
| --- | --- | --- | --- | --- | --- |
| 042-501 | Cesarean section | male | 160 | 0.442 | Dam |
| 042-502 | Cesarean section | female | 160 | 0.478 | Dam |
| 042-504 | Cesarean section | female | 160 | 0.5 | Dam |
| 042-505 | Cesarean section | female | 161 | 0.507 | Dam |
| 042-506 | Cesarean section | female | 160 | 0.445 | Dam |
| 042-507 | Cesarean section | female | 160 | 0.553 | Dam |
| 042-508 | Cesarean section | male | 160 | 0.5 | Peer |
| 044-501 | Cesarean section | male | 160 | 0.624 | Dam |
| 044-502 | Cesarean section | male | 160 | 0.498 | Peer |
| 044-503 | Cesarean section | male | 159 | 0.519 | Dam |
| 044-504 | Cesarean section | female | 159 | 0.465 | Dam |
| 044-505 | Cesarean section | female | 161 | 0.434 | Dam |
| 044-506 | Cesarean section | female | 160 | 0.563 | Peer |
| 044-507 | Cesarean section | female | 163 | 0.5 | Dam |
| 044-508 | Natural | female | 159 | 0.44 | Dam |
| 044-509 | Natural | male | 160 | 0.54 | Dam |
| 044-510 | Cesarean section | male | 160 | 0.594 | Dam |
| 044-511 | Cesarean section | female | 160 | 0.6 | Dam |
| 044-512 | Cesarean section | female | 159 | 0.44 | Dam |
| 044-513 | Cesarean section | female | 160 | 0.534 | Dam |
| 044-514 | Cesarean section | male | 160 | 0.442 | Dam |
| 044-515 | Cesarean section | female | 159 | 0.468 | Dam |
| 044-516 | Cesarean section | male | 160 | 0.6 | Dam |
| 044-517 | Cesarean section | male | 159 | 0.479 | Dam |
| 044-518 | Cesarean section | male | 155 | 0.413 | Dam |
| 044-520 | Cesarean section | female | 159 | 0.459 | Dam |
| 044-522 | Cesarean section | male | 158 | 0.501 | Dam |
| 044-523 | Cesarean section | female | 158 | 0.515 | Dam |
| 044-524 | Cesarean section | female | 160 | 0.504 | Peer |
| 044-525 | Cesarean section | female | 160 | 0.484 | Peer |
| 044-526 | Cesarean section | male | 160 | 0.486 | Dam |
| 044-527 | Cesarean section | male | 159 | 0.505 | Dam |
| 044-528 | Cesarean section | female | 158 | 0.561 | Dam |
| 044-530 | Cesarean section | male | 160 | 0.551 | Dam |
| 044-531 | Natural | male | 159 | 0.556 | Dam |
| 044-532 | Cesarean section | male | 160 | 0.601 | Dam |
| 044-533 | Cesarean section | male | 162 | 0.59 | Dam |
| 046-501 | Cesarean section | female | 160 | 0.493 | Dam |
| 046-502 | Cesarean section | female | 160 | 0.536 | Dam |
| 046-505 | Cesarean section | female | 159 | 0.55 | Dam |
| 046-506 | Cesarean section | male | 158 | 0.558 | Peer |

**Supplemental Table 12.** Infant behavior definitions

| **Infant behavior** | **Definition** |
| --- | --- |
| Mother-infant home cage observations | |
| Mutual ventral contact | Percent of time spent in ventral-ventral contact between dam and infant. |
| Nipple contact | Percent of time spent in oral contact by the infant with the dam’s nipple. |
| Proximity to mother (Together) | Percent of time dam and infant spent in physical contact or within monkey arm’s length of each other. |
| Locomotion | Percent of time spent in any self-induced change in location, including. Includes walking, climbing, rolling, hopping, bouncing, and dropping from ceiling to the floor of the home cage. |
| Puzzle Feeder | |
| Level Complete | Drop of the treat to the floor of the puzzle feeder where the treat can be retrieved |
| Individual attempt | Insertion of a finger into the hole nearest the food treat |
| Digit isolation | Identification of the number of digits the infant was using to manipulate the treat in the puzzle feeder |
| Motor coordination | Overall assessment of motor coordination during the task |
| Fine Motor PVC Pipe Test | |
| Oral exploration | % of time infant was orally exploring the PVC pipe |
| Frosting exploration | % of time infant was exploring the frosting from the PVC pipe |
| Digit isolation | % of time infant was using individual digits to manipulate frosting or raisins in our outside the PVC pipe |
| Sensory Response | |
| Infant approach | Dichotomous response of yes or no if the infant approached the sensory stimuli |
| Infant time to approach | Categorically recorded as >2 minutes, 1-2 minutes, or <1 minute |
| Visual approach | Infant moved toward the sensory stimulus at a distance of within one body length |
| Touch approach | Infant directly touched the sensory stimulus. Not only the metal rod. |
